# Bacteria-induced activation of a fungal silent gene cluster is controlled by histone deacetylase Sirtuin E

**DOI:** 10.1101/2023.12.05.569573

**Authors:** Nils Jäger, Maira Rosin, Maria C. Stroe, Axel A. Brakhage, Thorsten Heinzel

## Abstract

Filamentous fungi encode an untapped reservoir of natural products whose biosynthesis enzymes are often encoded by gene clusters. The majority of these gene clusters are only activated under distinct environmental conditions such as the presence of distinct neighbouring microorganisms but not under standard laboratory conditions. Previously, we provided evidence for such a scenario with the specific activation of the silent *ors* gene cluster in the filamentous fungus *Aspergillus nidulans* by the bacterium *Streptomyces rapamycinicus*. The bacterium triggered the activation of the GcnE histone acetyltransferase that acetylated histone 3 in nucleosomes of the *ors* gene cluster and the *basR* transcription factor, and thereby the gene cluster. The inducing compound was shown to be the bacterial arginoketide azalomycin F. Here, by inhibitor studies with the pan-sirtuin inhibitor nicotinamide (NAM) the involvement of a sirtuin HDAC was implied. Accordingly, deletion of all six putative sirtuin-encoding genes (*sirA*-*E* and *hstA*) revealed that only deletion of *sirE* led to production of orsellinic acid by *A. nidulans* without the need of the bacterium. Also other effects on growth and colony morphology due to NAM were phenocopied by the *sirE* deletion mutant. Addition of NAM did not compensate for the loss of the BasR transcription factor required for activation of the *ors* gene cluster. Collectively, SirE is the negative regulator of the bacteria-induced activation of the *ors* BGC. In line, addition of NAM to monocultures of *Aspergillus mulundensis* encoding a sirtuin E with highest similarity to the *A. nidulans* protein also activated the *ors* BGC in this fungus.

## Introduction

Filamentous fungi are important producers of low molecular mass natural products (NPs) many of which have found pharmaceutical and biotechnological applications (Yu and Keller, 2005). Prominent examples are the antibiotics cephalosporin C and penicillin G, the immunosuppressant cyclosporine A or the antimycotic substance griseofulvin (Oxford *et al*., 1939; Burton and Abraham, 1951; Zocher *et al*., 1986; Brakhage, 2013). The genes required for their biosynthesis are often physically linked in biosynthetic gene clusters (BGCs). Analyses of sequenced fungal genomes by search programs such as SMURF or AntiSMASH allow to identify BGCs (Khaldi *et al*., 2010; Medema *et al*., 2011). Based on such analyses, the model filamentous fungus *Aspergillus nidulans* contains at least 66 putative BGCs, most of which are not yet assigned to NPs (Romsdahl and Wang, 2019). This is due to the observation that many BGCs remain inactive or are lowly expressed under standard laboratory conditions (Brakhage, 2013). They can be only activated when the correct environmental stimulus is provided that can consist of a microbial partner (Schroeckh *et al*., 2009; Hotter *et al*., 2021; Netzker *et al*., 2015).

An early example is the activation of the cryptic *ors* gene cluster of *A. nidulans* by co-cultivation of the fungus with the soil-dwelling bacterium *Streptomyces rapamycinicus* and its closest relative *S. iranensis* (Nützmann *et al*., 2011; Krespach *et al*., 2023). Recently, we identified a group of polyketides named arginoketides that represent the bacterial signal inducing the silent *ors* gene cluster (Krespach *et al*., 2023). It was particularly interesting to notice that *S. rapamycinicus* was able to re-program the fungal histone modification machinery (Nützmann *et al*., 2011; Fischer *et al*., 2018; Netzker *et al*., 2018). Increased acetylation of histone H3 lysine (K) 9 and H3K14 of nucleosomes present in orsellinic acid biosynthesis gene promoters was detected upon co-cultivation and was shown to be required for activation of the *ors* gene cluster. Also a number of nucleosomes in promoters of transcription factors with increased histone acetylation were detected leading to the discovery of the transcription factor BasR that is required for the activation of the *ors* gene cluster. The responsible histone acetyltransferase (HAT) was found to be GcnE, a member of the SAGA/ADA complex (Nützmann *et al*., 2011). Upon interaction of the fungus with the bacterium, GcnE acetylates histone H3 leading to the expression of the *ors* biosynthesis genes and production of orsellinic acid and derivatives. The Myb-like transcriptional regulator BasR represents the major fungal regulatory node transducing the bacterial signal to activation of the *ors* gene cluster (Fischer *et al*., 2018). Accordingly, an important question concerns the reversion of the acetylation signal, *i*.*e*., how the activation is terminated. For deacetylation of histones, HAT-counteracting histone deacetylases (HDACs) are required. By inhibitor studies, gene deletion analyses and knockdown studies we identified the silent information regulator SirE (Itoh *et al*., 2017) as the HDAC terminating the induction of the *ors* BGC by *S. rapamycinicus*. SirE is the negative regulator of the bacteria-induced *ors* BGC.

## Results

### Pan-sirtuin inhibitor nicotinamide (NAM) affects radial growth, conidiation and natural product formation of *A. nidulans*

Analysis of the literature and our experiments with available deletion mutants of HDACs, *i*.*e*., HosA, HosB, RpdA and HdaA (Graessle *et al*., 2000) indicated that these HDACs are not involved in the regulation of the *ors* gene cluster because the production of orsellinic acid induced by *S. iranensis* was still fully detectable in co-cultures with the respective mutants (data not shown). Therefore, we also considered NAD^+^-dependent class III HDACs known as sirtuins which are encoded by six genes in *A. nidulans* designated *sirA, sirB, sirC, sirD, sirE*, and *hstA* (Shwab *et al*., 2007; Itoh *et al*., 2017a, 2017b). The catalytic core of sirtuins is composed of a Zn^2+^-binding domain and a large NAD^+^-binding domain containing a Rossmann fold (Min *et al*., 2001). In contrast to class I and II HDACs, sirtuins require NAD^+^ as a co-factor for their catalytic activity (Sauve *et al*., 2001). During catalysis, NAD^+^ is hydrolyzed to NAM while a deacetylated peptide and both 2’- and 3’-O-acetyl-ADP-ribose are formed (Sauve *et al*., 2001). Hence, NAM is a known non-competitive inhibitor of sirtuin catalytic activity (Sauve *et al*., 2001). In contrast to artificial sirtuin inhibitors, the physiological compound NAM can be metabolized to NAD^+^ explaining the need for higher concentrations (Imai and Guarente, 2016).

To investigate whether sirtuins are involved to terminate the bacterial azalomycin F signal inducing the *ors* gene cluster, NAM was applied in concentrations which are in line with published data (Avalos *et al*., 2005; Wurtele *et al*., 2010; Stevenson and Liu, 2011; Griffin *et al*., 2017). The supplementation of *A. nidulans* strain A1153 with 10-40 mM NAM significantly reduced the radial growth and conidia formation of colonies on AMM agar plates (Figure S1). The cytotoxicity of NAM was already shown before for the fungus *Candida albicans*, in which repression of HAT HST3 by NAM led to RTT109-dependent H3K56 hyperacetylation (Wurtele *et al*., 2010). In line, in *A. fumigatus* the loss of the orthologous HAT RTT109 led to decreased H3K56- and H3K9 acetylation (Zhang *et al*., 2020). To determine the sensitivity of *A. nidulans* to NAM, knockout strains of *RTT109* and *gcnE* were compared with the wild type. As shown for *C. albicans*, the loss of *RTT109* led to higher tolerance against NAM treatment (Figure S2), also suggesting counteracting functions of *A. nidulans* sirtuins and RTT109.

Supplementation of *A. nidulans* with NAM led to enhanced production and secretion of a brownish metabolite around the marginal zones of the colony (Figure S1A). Therefore, extracts of NAM-treated cultures were analyzed by LC-MS. They contained orsellinic acid, lecanoric acid and F-9775A, F-9775B compounds produced by the *ors* BGC without the need of the bacterium (Figure 1A). To relate the compounds to expression of *ors* genes we applied an *A. nidulans orsA*p-*nluc*-GFPs reporter strain (Krespach *et al*., 2023) (Figure 1C). Upon treatment of this strain with NAM both increased nanoluciferase activity (Figure 1D) and GFPs-derived fluorescence were detected (Figure 1E). This finding indicates enhanced promoter activity of the *orsA* promoter upon addition of NAM.

**Figure 1.**
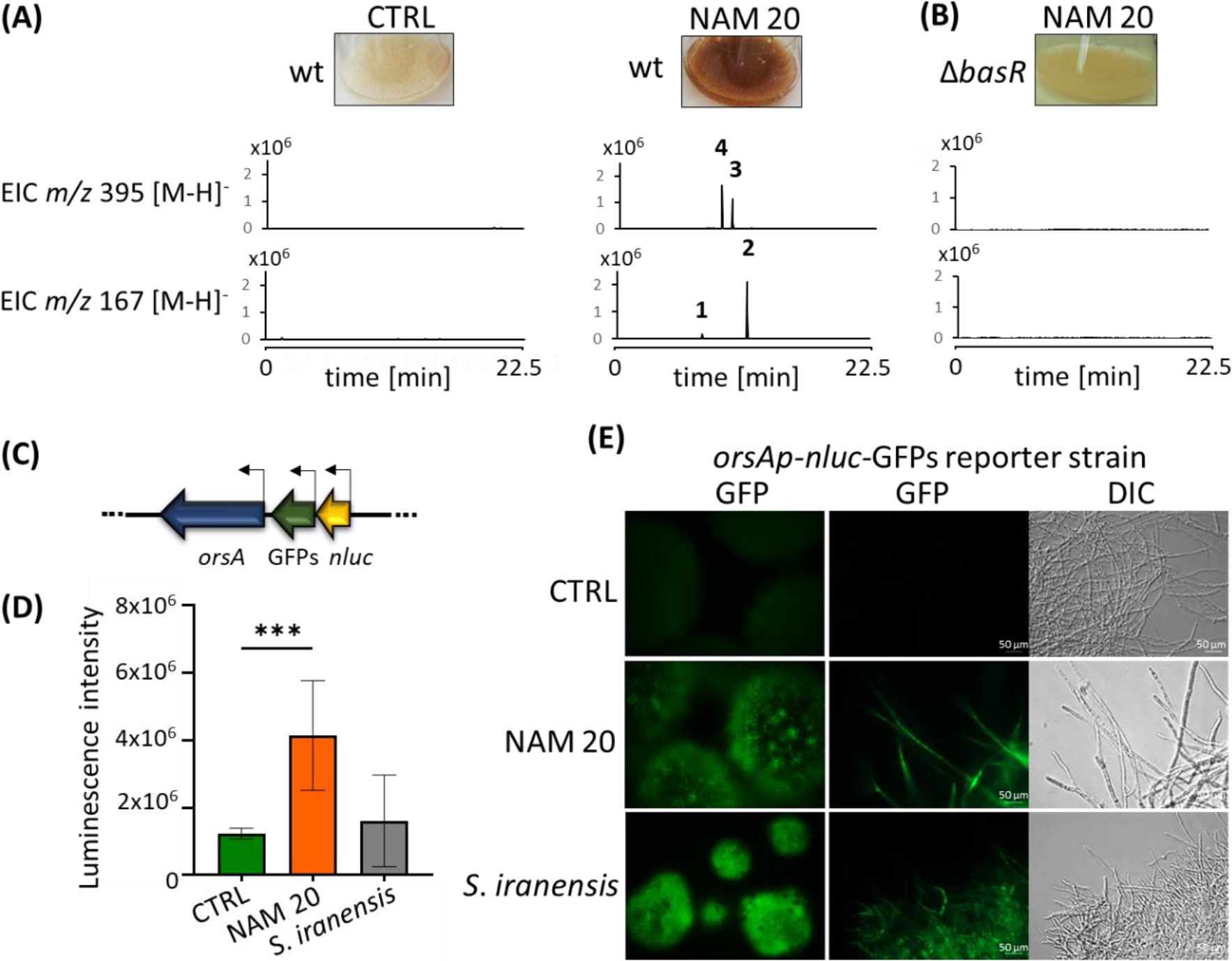
BasR-dependent production of *ors* BGC-derived NPs during NAM treatment of *A. nidulans*. **(A)** *A. nidulans* colonies grown in AMM with or without supplementation of 20 mM NAM for 24 h at 37°C with 200 rpm (left). Pictures of cultures and extracted ion chromatograms (EICs) are shown for m/z 167 [M-H]^-^ and m/z 395 [M-H]^-^, corresponding to *ors* gene cluster-derived NPs. Marked peaks correspond to orsellinate (1), lecanorate (2), F-9775A (3) and F-9775B (4). **(B)** Treatment of a *basR* deletion strain with 20 mM NAM. **(C)** Schematic of genomic organization of *A. nidulans orsA*p-nLuc-GFPs reporter strain. **(D)** Luminescence intensities of the *A. nidulans orsA*p-nLuc-GFPs reporter strain grown with 20 mM NAM, co-cultivated with *S. iranensis*, or with PBS (CTRL). Bars represent mean ± SD. Significance was assessed by an unpaired *t*-test (** *p* < 0.02; *** *p* < 0.002). **(E)** Fluorescence images of *A. nidulans orsA*p-nLuc-GFPs reporter strain grown with 20 mM NAM, co-cultivated with *S. iranensis* or with PBS (CTRL Scale bar: 50 μm).

Since the Myb-like transcription factor (TF) BasR was shown to be also essential for the activation of the *ors* BGC during co-cultivation of *A. nidulans* with *S. rapamycinicus* (Fischer *et al*., 2018), we investigated whether NAM-induced changes in histone modifications compensate for the lack of *basR* in activating the *ors* BGC. As shown in Figure 1B, this was not the case, BasR was required for the NAM effect. To investigate, whether the effects of NAM on conidiation and radial growth also depend on BasR, a *basR* deletion strain was grown on AMM agar plates in presence of 10-40 mM NAM. In the Δ*basR* mutant strain, there was no difference in the analyzed phenotypes compared to the wild type when treated with NAM suggesting the involvement of other transcriptional regulators than BasR for these phenotypes (Figure S2). Although no *ors* gene cluster-derived NPs were detected by LC-MS in a *basR* deletion strain (Figure 1B), the presence of a yellowish compound around the marginal zones of the colony indicated the potential activation of other BGCs by NAM-supplementation (Figure S2).

### Morphological and physiological changes due to NAM are mirrored by the phenotype of Sirtuin E inhibition

By BLASTp analyses based on the amino acid sequence of SIR2 of *S. cerevisiae*, in the genome of the *A. nidulans* strain FGSC A4 six putative sirtuin orthologous proteins were identified. The deduced amino acid sequences of all sirtuins of *S. cerevisiae, Homo sapiens*, and *A. nidulans* were aligned using MUSCLE and a phylogenetic tree was constructed using MEGA6 (Figure S3). The resulting phylogenetic tree and the identified SIR2 homologs were supported by existing literature (Kawauchi *et al*., 2013). Nuclear localization of sirtuins is a prerequisite for deacetylation of histones. According to the program NLS Mapper, only SirE and SirA exhibit nuclear localization sequences (NLSs) (Figure S3B). This finding agrees with results obtained by fluorescence microscopy with GFP-tagged SirA and SirE gene fusions (Itoh *et al*., 2017a).

To identify the sirtuin responsible for termination of the bacteria-induced activation of the *ors* BGC and for the effects observed upon NAM-treatment, we deleted all six putative sirtuin-encoding genes of *A. nidulans* (Figure S4A-F). Colonies of knockout strains of *sirA* (*AN10449*), *sirB* (*AN7461*), *sirC* (*AN1782), sirD* (*AN7461*) and *hstA* (*AN11067*) did not reveal an aberrant morphology in comparison to the wild type (Figure 2A). Furthermore, the resistance to menadione and Congo red inducing reactive oxygen species and cell wall disturbance, respectively, was not affected by loss of the above mentioned sirtuins (Figure S5). This was different for the *sirE* (*AN1226*) deletion strain that showed strongly reduced radial growth and conidiation, as previously reported (Itoh *et al*., 2017a). Interestingly, the *sirE* mutant strain clearly produced orsellinic acid without the need of bacteria, as also seen for the wild type treated with NAM (Figure 2B). By contrast, none of the other sirtuin deletion mutant strains produced orsellinic acid or derivatives thereof (Figure 2C).

**Figure 2.**
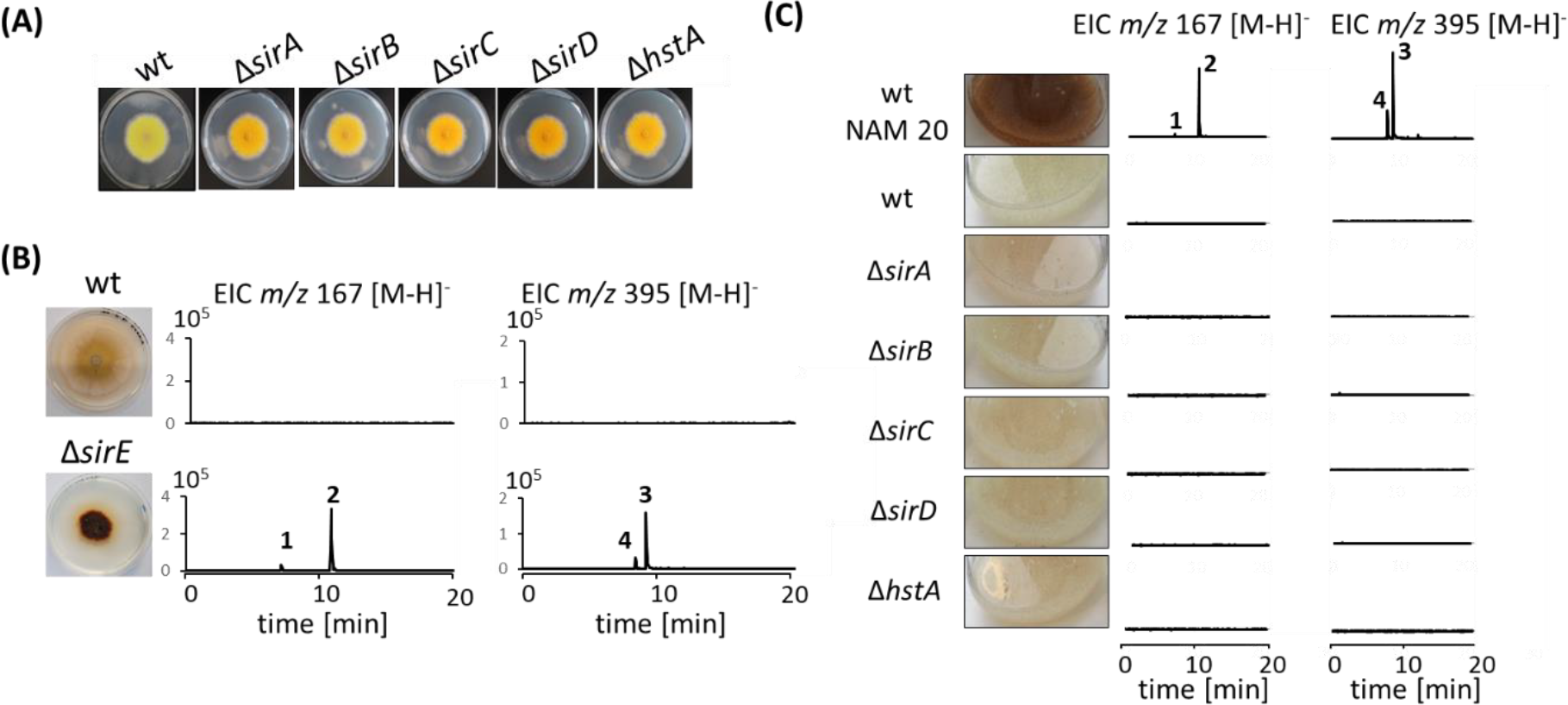
Loss of SirE affects biosynthesis of *ors* compounds and colony morphology. **(A)** *A. nidulans* colonies of wt, Δ*sirA*, Δ*sirB*, Δ*sirC*, Δ*sirD*, and Δ*hstA* mutant strains grown on AMM agar. Conidia (10^5^) of indicated *A. nidulans* strains were point-inoculated on AMM agar plates and cultivated for 3 d at 37 °C. **(B)** Colony of wt and Δ*sirE* mutant strain on AMM agar plates grown for 7 d at 37 °C. LC-MS profiles of EICs for m/z 167 [M-H]^-^ and m/z 395 [M-H]^-^. Marked peaks correspond to orsellinate (1), lecanorate (2), F-9775A (3), and F-9775B (4). **(C)** Liquid cultures of wt, Δ*sirA*, Δ*sirB*, Δ*sirC*, Δ*sirD*, and Δ*hstA* strains grown in AMM at 37°C with 200 rpm for 24 h. The wt strain treated with 20 mM NAM served as a positive control. LC-MS profiles of EICs for m/z 167 [M-H]^-^ and m/z 395 [M-H]^-^. Marked peaks correspond to orsellinate (1), lecanorate (2), F-9775A (3), and F-9775B (4).

### Downregulation of SirE leads to the activation of the *ors* gene cluster

As further experiments were hampered by the drastic phenotype of the *sirE* deletion strain, we generated a strain with an exchange of the native *sirE* promoter by the regulatable *xylP* promoter of the β-1,4-endoxylanase gene from *Penicillium chrysogenum* (Zadra *et al*.,

2000). This promoter has been successfully used for downregulation of essential genes such as the class II HDAC RpdA in *A. nidulans* and *A. fumigatus* (Tribus *et al*., 2010; Bauer *et al*., 2019). While xylose is required to induce the expression of *xylP*p-regulated genes, glucose acts as a repressor in the absence of xylose (Figure S6A). However, as the strain did not show a growth defect even under *sirE* repressing conditions with glucose, the knockdown efficiency of the *xylP*p-*sirE* strain was apparently low under the tested conditions indicating some degree of leakiness of the *xylP* promoter (Figure S7), as reported before (Bauer *et al*., 2016; Misslinger *et al*., 2019). The general expression of *sirE* might be low in the wild type under the tested conditions because Itoh *et al*. detected enhanced expression during the stationary phase. Thus, low levels of SirE protein seem to be sufficient to silence SirE-regulated genes effectively. In line, analysis of the mRNA steady-state level by qRT-PCR showed that the slight repression of the *sirE* gene when under control of *xylP*p was sufficient to allow expression of *basR* and likely as a result, of *orsA* and *orsD* (Figure 3C-D). These findings agree well with our previous results on the *basR* promoter that showed higher acetylation levels in response to co-cultivation with bacteria (Fischer *et al*., 2018) and this mark is most likely removed by SirE. Our expression analysis also confirmed that overexpression of *sirE* due to the presence of xylose in the medium was very effective (Figure 3A). High levels of SirE apparently affected the overall colony morphology leading to ragged edges of the colony (Figure S7). LC-MS^2^ analysis confirmed that a slight repression of *sirE* was sufficient to induce the production of orsellinic acid, lecanoric acid, and compounds F-9775A and F-9775B (Figure 3B). It is worth to notice that the overexpression of *sirE* by xylose did not completely abolish the production of orsellinic and lecanoric acid when grown with *S. rapamycinicus* or arginoketides as seen in a *basR* deletion strain (Figure S8). Together, these findings indicate that SirE acts as a repressor of the *ors* BGC during vegetative growth likely by removing acetylation marks of H3K9 and K14.

**Figure 3.**
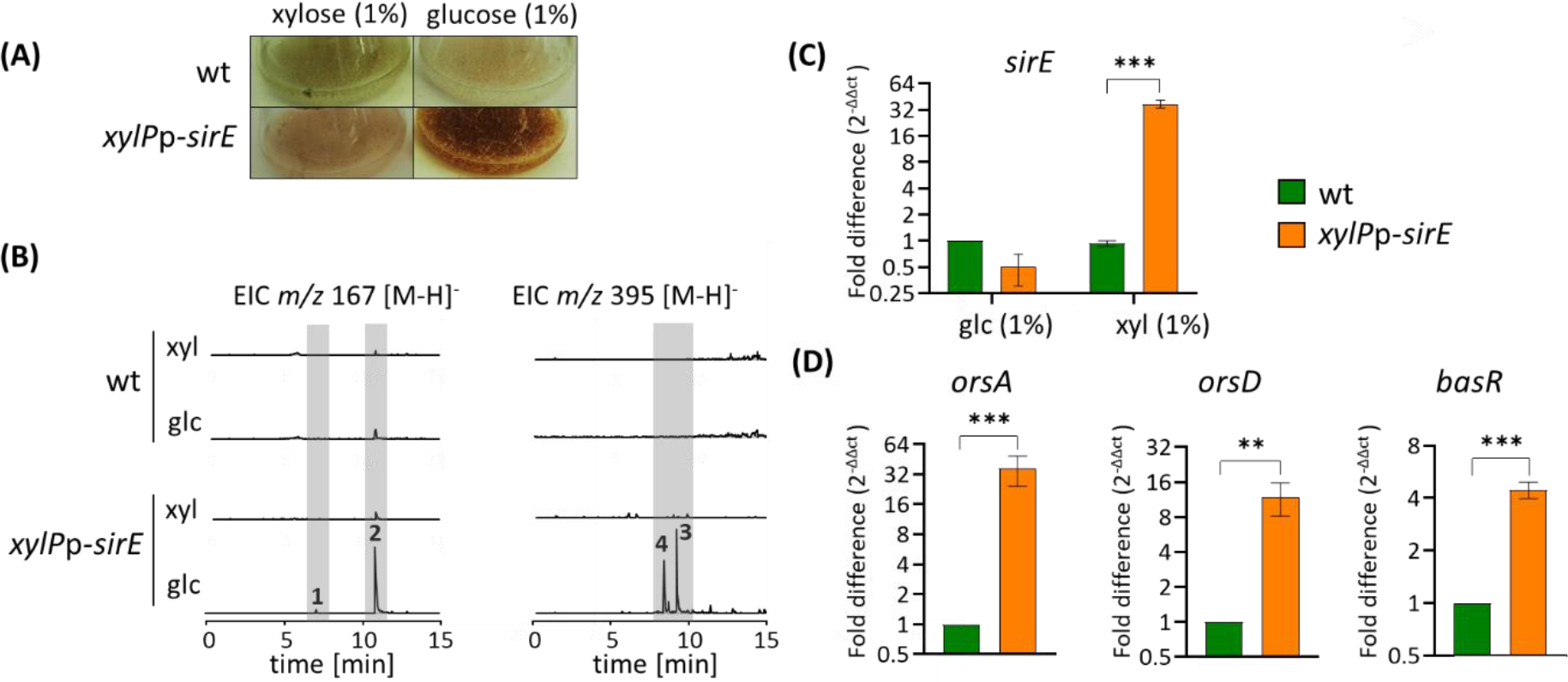
Growth under *xylP*p-repressing conditions activates the *ors* gene cluster in the *xylP*p-*sirE* strain. **(A)** Cultures of *A. nidulans* wt and *xyl*p-*sirE* strain grown in AMM in mono-culture under repressing (1 % (w/v) glc) or inducing (1 % (w/v) xyl) conditions for 24 h at 37 °C. **(B)** LC-MS profiles of EICs are depicted for m/z 167 [M-H]^-^ and m/z 395 [M-H]^-^. Marked peaks correspond to orsellinate (1), lecanorate (2), F-9775A (3), and F-9775B (4). Relative transcript levels for **(C)** *sirE* under inducing and repressing conditions, **(D)** *orsA, orsD*, and *basR* were obtained by qRT-PCR of the *xylP*p-*sirE* strain grown under repressing conditions (for *orsA, orsD* and *basR*) with 1 % (w/v) glucose. Data were normalized against β-actin (*AN6542*) according to the ΔΔCt method (n=3, bars stand for mean ± SD). Significance was assessed by an unpaired *t*-test (** *p* < 0.02; *** *p* < 0.002).

### SirE is conserved among aspergilli

To further investigate the presence of SirE in distantly related aspergilli, we compared SirE among different aspergilli by BLASTp analysis based on the amino acid sequence of SirE. The use of the program MUSCLE allowed us to generate a phylogenetic tree (Altschul *et al*., 1990; Tamura *et al*., 2013). Orthologs of SirE were identified in all genomes of the analyzed *Aspergillus* sections. Notably, three *Aspergillus spp*. which are known to produce SMs during the interaction with *S. rapamycinicus*, namely *A. sydowii, A. versicolor* and *A. fumigatus*, encode *sirE* homologs (Figure 4). SirE of the *A. mulundensis* strain DSM5745_06405 showed the highest similarity to SirE of *A. nidulans* (Figure 4). Since this strain was not available we analyzed the genome of *A. mulundensis* strain CBS140610 by antiSMASH (Medema *et al*., 2011) and BLASTp (Altschul *et al*., 1990) which revealed the presence of a putative *ors* gene cluster, as well as a the presence of a BasR-homologous protein, respectively (Figure S9). According to these analyses, the putative *ors* gene cluster of *A. mulundensis* consists of a putative *orsA* (DSM5745_00010), *orsB* (DSM5745_00008), *orsD* (DSM5745_00006) and *orsE* (DSM5745_00005) gene and is located in a sub-telomeric region (Figure S9). Furthermore, the genome of *A. mulundensis* encodes a R3-Myb-like transcription factor with a high similarity to BasR of *A. nidulans*. Likewise, *A. mulundensis* displayed a similar phenotype as seen for *A. nidulans* when malt agar was supplemented with 10-40 mM of NAM (Figure 5A). The colony radial growth as well as the conidiation were significantly reduced (Figure 5B). Similar to *A. nidulans*, orsellinic acid was identified in an *A. mulundensis* co-culture with *S. rapamycinicus* as well as after the addition of 40 mM NAM (Figure 5). This further confirms the conserved function of sirtuins in aspergilli.

**Figure 4.**
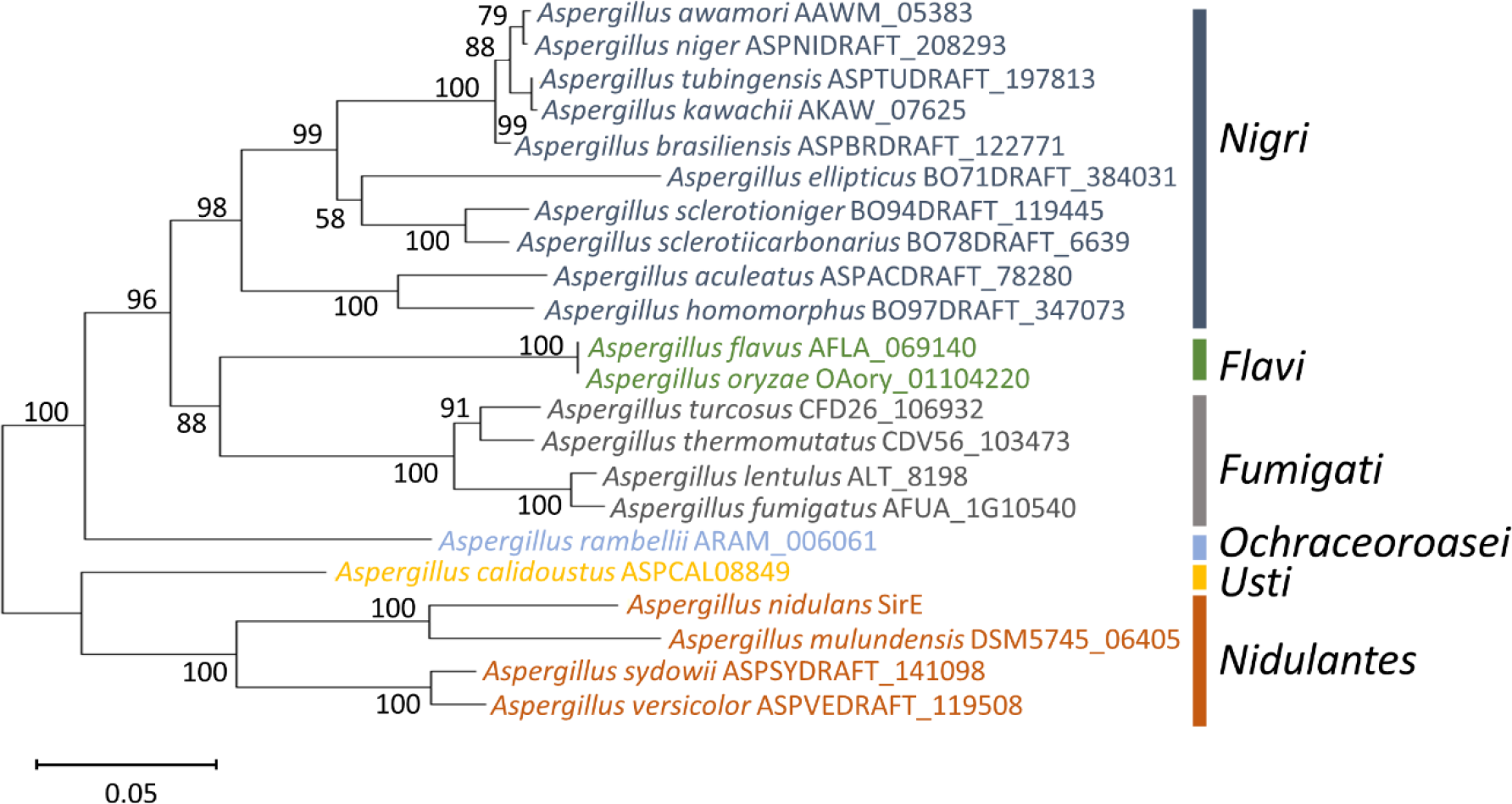
Phylogenetic tree of SirE orthologs of different aspergilli based on amino acid sequences. MUSCLE alignment and neighbour-joining were performed with 1000 bootstrap replicates of the listed *Aspergillus* strains and the amino acid sequences of their corresponding SirE homologs in MEGA6 (Edgar, 2004; Tamura *et al*., 2013). The percentage of replicate trees in which the associated taxa cluster, is shown next to the branches.

**Figure 5.**
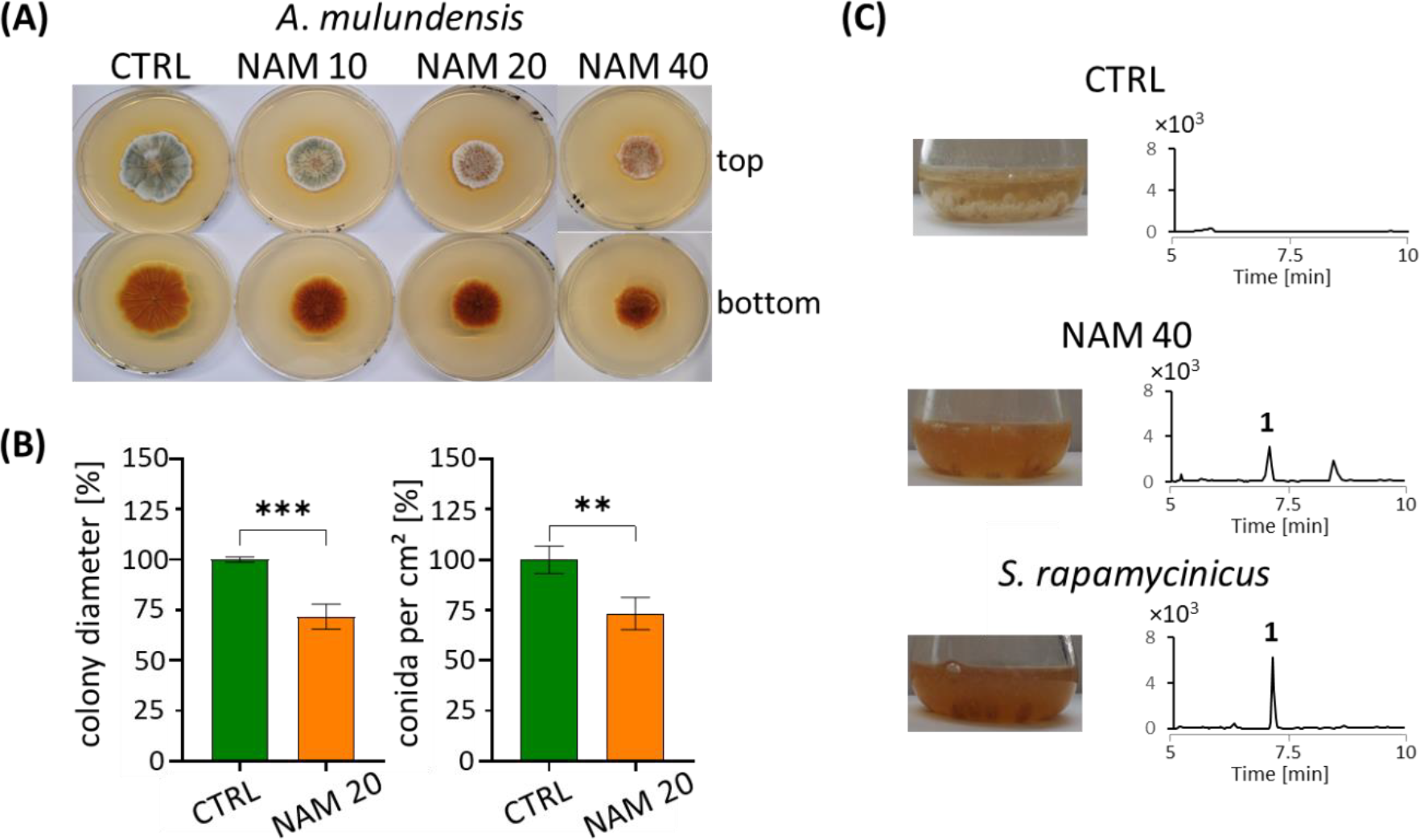
Phenotypic analysis of NAM-treated *A. mulundensis*. **(A)** Colonies on agar plates shown from top and bottom. Conidia (10^6^) of *A. mulundensis* were point-inoculated on malt agar plates supplemented with and without 10-40 mM NAM and grown for 7 d at 26 °C. **(B)** Colony diameter and number of produced conidia in response to NAM (n=3, bars stand for mean ± SD). Significance was assessed by an unpaired *t*-test (** *p* < 0.02; *** *p* < 0.002). **(C)** Pictures of liquid (co-)cultures from *A. mulundensis* after 42 h of growth at 26 °C. (left) and EICs from culture extracts are depicted for m/z 167 [M-H]^-^ (right). Marked peaks correspond to orsellinate (1).

## Discussion

Advances in sequencing of fungal genomes and their analysis have shown that many fungal BGCs remain silent under standard laboratory conditions (Brakhage, 2013). In recent years environmental stimuli for their induction have been uncovered as well as the importance of chromatin modifiers in activating silent BGCs (Nützmann *et al*., 2011; Krespach *et al*., 2023). For example, loss of CclA of the H3K4 methylating COMPASS complex led to the appearance of *ors*-derived compounds and gliotoxin in culture media of *A. nidulans* and *Aspergillus fumigatus*, respectively (Bok *et al*., 2009; Palmer *et al*., 2013). Likewise, the methyltransferase LaeA and the HAT GcnE were shown to be master regulators of NP biosyntheses and to affect the expression of different BGCs of filamentous fungi (Nützmann *et al*., 2011; Kim *et al*., 2013; Rösler *et al*., 2016; Martín, 2017). An interesting aspect concerning the ecology and regulation is the bacteria-induced acetylation of the histone marks H3K9 and H3K14 that was shown to be pivotal for the activation of the *ors* BGC and other BGCs in *A. nidulans* (Nützmann *et al*., 2013). HAT-counteracting HDACs catalyze the removal of acetyl groups from lysine residues from histone and non-histone proteins and are thus required to silence previously activated genes. Until today, such an HDAC that counteracts bacteria-induced BGC activation had not been identified. Here, we discovered that sirtuin E is the sought-after HDAC.

Deacetylation of histone residues leads to formation of compact heterochromatin, which is not accessible for the transcriptional machinery (Workman and Kingston, 1998). Thus, deacetylation of histones is generally regarded as a key mechanism in silencing of transcription (Workman and Kingston, 1998). Consequently, the use of HDAC inhibitors has been shown to be a successful method to induce otherwise silent BGCs. For example, the use of the classical HDACi SAHA was shown to enhance the production of a complex series of perylenequinones in *Cladosporium cladosporioides* (Williams *et al*., 2008) and to activate the biosynthesis of the cryptic NP BGC for nygerone A in *A. niger* and the *ors* gene cluster in *A. nidulans* (Henrikson *et al*., 2009; Nützmann *et al*., 2011).

Besides classical Zn^2+^-dependent HDACs, fungal genomes encode NAD^+^-dependent class III HDACs known as sirtuins which were shown to regulate cryptic BGCs in filamentous fungi. For example, NAM-treatment of the entomopathogenic fungus *Chaetomium mollipilium* and of the endophytic fungus *Graphiopsis chlorocephala* led to the discovery of the novel C_13_-polyketides mollipilin A-E and the benzophenones cephalanones A-F (Asai *et al*., 2012; Chung *et al*., 2013). Similarly, *Chaetomium cancroideum* produced previously not observed NPs upon NAM treatment (Asai *et al*., 2016). Here, we show that growth of *A. nidulans* in presence of NAM led to reduced radial growth and conidiation as well as to the specific activation of the cryptic *ors* BGC. The additional phenotypes to the activation of the NP formation suggest a major effect of NAM on different metabolic pathways and might also explain the antifungal activity of NAM (Wurtele *et al*., 2010). As shown here, all the effects of NAM-supplementation were phenocopied by deletion and mostly by repression of the sirtuin SirE and not by deletion of any other of the six sirtuin genes. Similarly, loss of the SirE ortholog HstD affected the conidiation of *A. oryzae* (Kawauchi *et al*., 2013).

SirE was previously shown to specifically regulate H3K9 and H3K56 deacetylation in *A. nidulans* (Itoh *et al*., 2017a). In line, the reduction of the SirE orthologous Hst3 in *C. albicans* was shown to affect H3K56 deacetylation, and deletion of the gene was lethal (Wurtele *et al*., 2010). Here, overexpression as well as the deletion of *sirE* showed detrimental effects on the morphology of *A. nidulans*. Moreover, the loss of the HAT RTT109 showed higher resistance to high concentrations of NAM. Together, these findings suggest that a tight balance between the counteracting catalytic activities of RTT109 and SirE is essential. In *S. cerevisiae*, the knockout of the *sirE* orthologs *hst3*/*hst4* led to loss of telomeric silencing (Aparicio *et al*., 1991; Brachmann *et al*., 1995). In particular, the *hst3*/*hst4*-deficient strains accumulated acetylation of H3 at K56 on newly synthesized histones by the HAT RTT109 (Yang *et al*., 2008). Follow-up studies demonstrated that deacetylation of histone H3K56 by Hst3 and Hst4 was important to minimize genomic instability during replication (Hachinohe *et al*., 2011). This finding is in line with data from *C. albicans*, where the knockdown of the *sirE* ortholog *hst3* displayed enhanced H3 acetylation at K56 by RTT109 (Wurtele *et al*., 2010). Similarly, the human SirE ortholog SIRT6 was shown to deacetylate the histone marks H3K9 and H3K56 at telomeric ends (Michishita *et al*., 2009; Tennen *et al*., 2011), as also shown for the SirE ortholog Nst-3 of *Neurospora crassa* that specifically silenced genes located near the telomeric ends (Smith *et al*., 2008). It is therefore not surprising that SirE of *A. nidulans* might be involved in the regulation of sub-telomerically located genes such as the *ors* gene cluster. This finding also underlines the importance of SirE for finding novel bioactive molecules as fungal BGCs are often located sub-telomerically and are silenced due to the spread of facultative heterochromatin (Palmer and Keller, 2010). Due to its dependence on NAD^+^, SirE could also play a role in linking the expression of BGCs to the metabolic state of the cell. Besides its effects on histone proteins, SirE might influence the acetylation status of non-histone proteins, as shown for the human HAT Gcn5. Its deacetylation by SIRT6 affected its catalytic activity (Dominy *et al*., 2012). It is conceivable that co-cultivation of *A. nidulans* with *S. rapamycinicus* and its produced compound azalomycin F influence SirE that either acts directly on GcnE or on the acetylation status of the *basR* and *ors* gene promoters, thereby regulating the expression of the *ors* BGC.

## Acknowledgements

We thank Christina Täumer und Jonas Fiedler for excellent technical assistance. We also thank Dr. Stefan Graessle for kindly providing HDAC mutant strains. Financial support by the DFG-funded Collaborative Research Center 1127 *ChemBioSys* (project B02 No. 239748522) and the Cluster of Excellence Balance of the Microverse under Germany’s Excellence Strategy - EXC 2051 - Project-ID 390713860 is gratefully acknowledged.

## References

Albright, J.C., Henke, M.T., Soukup, A.A., McClure, R.A., Thomson, R.J., Keller, N.P., Kelleher, N.L., 2015. Large-Scale Metabolomics Reveals a Complex Response of Aspergillus Nidulans to Epigenetic Perturbation. ACS Chem. Biol. 10, 1535–1541.

Altschul, S.F., Gish, W., Miller, W., Myers, E.W., Lipman, D.J., 1990. Basic Local Alignment Search Tool. J. Mol. Biol. 215, 403–410.

Aparicio, O.M., Billington, B.L., Gottschling, D.E., 1991. Modifiers of Position Effect Are Shared between Telomeric and Silent Mating-Type Loci in S. Cerevisiae. Cell 66, 1279–1287.

Asai, T., Morita, S., Shirata, N., Taniguchi, T., Monde, K., Sakurai, H., Ozeki, T., Oshima, Y., 2012. Structural Diversity of New C13-Polyketides Produced by Chaetomium Mollipilium Cultivated in the Presence of a NAD+-Dependent Histone Deacetylase Inhibitor. Org. Lett. 14, 5456–5459.

Asai, T., Morita, S., Taniguchi, T., Monde, K., Oshima, Y., 2016. Epigenetic Stimulation of Polyketide Production in Chaetomium Cancroideum by an NAD+-Dependent HDAC Inhibitor. Org. Biomol. Chem. 14, 646–651.

Avalos, J.L., Bever, K.M., Wolberger, C., 2005. Mechanism of Sirtuin Inhibition by Nicotinamide: Altering the NAD + Cosubstrate Specificity of a Sir2 Enzyme. Mol. Cell 17, 855–868.

Bauer, I., Misslinger, M., Shadkchan, Y., Dietl, A.M., Petzer, V., Orasch, T., Abt, B., Graessle, S., Osherov, N., Haas, H., 2019. The Lysine Deacetylase RpdA Is Essential for Virulence in Aspergillus Fumigatus. Front. Microbiol. 10, 1–9.

Bauer, I., Varadarajan, D., Pidroni, A., Gross, S., Vergeiner, S., Faber, B., Hermann, M., Tribus, M., Brosch, G., Graessle, S., 2016. A Class 1 Histone Deacetylase with Potential as an Antifungal Target. MBio 7, 1–13.

Bok, J.W., Chiang, Y.M., Szewczyk, E., Reyes-Domingez, Y., Davidson, A.D., Sanchez, J.F., Lo, H.C., Watanabe, K., Strauss, J., Oakley, B.R., Wang, C.C.C., Keller, N.P., 2009. Chromatin-Level Regulation of Biosynthetic Gene Clusters. Nat. Chem. Biol. 5, 462–464.

Brachmann, C.B., Sherman, J.M., Devine, S.E., Cameron, E.E., Pillus, L., Boeke, J.D., 1995. The SIR2 Gene Family, Conserved from Bacteria to Humans, Functions in Silencing, Cell Cycle Progression, and Chromosome Stability. Genes Dev. 9, 2888–2902.

Brakhage, A.A., 2013. Regulation of Fungal Secondary Metabolism. Nat. Rev. Microbiol. 11, 21–32.

Brakhage, A.A., Van den Brulle, J., 1995. Use of Reporter Genes to Identify Recessive Trans-Acting Mutations Specifically Involved in the Regulation of Aspergillus Nidulans Penicillin Biosynthesis Genes. J. Bacteriol. 177, 2781–2788.

Burton, H.S., Abraham, E.P., 1951. Isolation of Antibiotics from a Species of Cephalosporium; Cephalosporins P1, P2, P3, P4, and P5. Biochem. J. 50, 168–174.

Chung, Y.M., Wei, C.K., Chuang, D.W., El-Shazly, M., Hsieh, C.T., Asai, T., Oshima, Y., Hsieh, T.J., Hwang, T.L., Wu, Y.C., Chang, F.R., 2013. An Epigenetic Modifier Enhances the Production of Anti-Diabetic and Anti-Inflammatory Sesquiterpenoids from Aspergillus Sydowii. Bioorganic Med. Chem. 21, 3866–3872.

Dominy, J.E., Lee, Y., Jedrychowski, M.P., Chim, H., Jurczak, M.J., Camporez, J.P., Ruan, H. Bin Feldman, J., Pierce, K., Mostoslavsky, R., Denu, J.M., Clish, C.B., Yang, X., Shulman, G.I., Gygi, S.P., Puigserver, P., 2012. The Deacetylase Sirt6 Activates the Acetyltransferase GCN5 and Suppresses Hepatic Gluconeogenesis. Mol. Cell 48, 900–913.

Fischer, J., Müller, S.Y., Netzker, T., Jäger, N., Gacek-Matthews, A., Scherlach, K., Stroe, M.C., García-Altares, M., Pezzini, F., Schoeler, H., Reichelt, M., Gershenzon, J., Krespach, M.K.C., Shelest, E., Schroeckh, V., Valiante, V., Heinzel, T., Hertweck, C., Strauss, J., Brakhage, A.A., 2018. Chromatin Mapping Identifies BasR, a Key Regulator of Bacteria-Triggered Production of Fungal Secondary Metabolites. Elife 7, 1–30.

Gibson, D.G., Young, L., Chuang, R.Y., Venter, J.C., Hutchison, C.A., Smith, H.O., 2009. Enzymatic Assembly of DNA Molecules up to Several Hundred Kilobases. Nat. Methods 6, 343–345.

Graessle, S., Dangl, M., Haas, H., Mair, K., Trojer, P., Brandtner, E.M., Walton, J.D., Loidl, P., Brosch, G., 2000. Characterization of Two Putative Histone Deacetylase Genes from Aspergillus Nidulans. Biochim. Biophys. Acta - Gene Struct. Expr. 1492, 120–126.

Griffin, S.M., Pickard, M.R., Orme, R.P., Hawkins, C.P., Williams, A.C., Fricker, R.A., 2017. Nicotinamide Alone Accelerates the Conversion of Mouse Embryonic Stem Cells into Mature Neuronal Populations. PLoS One 12, 1–17.

Hachinohe, M., Hanaoka, F., Masumoto, H., 2011. Hst3 and Hst4 Histone Deacetylases Regulate Replicative Lifespan by Preventing Genome Instability in Saccharomyces Cerevisiae. Genes to Cells 16, 467–477.

Henrikson, J.C., Hoover, A.R., Joyner, P.M., Cichewicz, R.H., 2009. A Chemical Epigenetics Approach for Engineering the in Situ Biosynthesis of a Cryptic Natural Product from Aspergillus Niger. Org. Biomol. Chem. 7, 435–438.

Hotter V., Zopf D., Kim H.J., Silge A., Schmitt M., Aiyar P., Fleck J., Matthäus C., Hniopek J., Yan Q., Loper J., Sasso S., Hertweck C., Popp J., Mittag M., 2021 A polyyne toxin produced by an antagonistic bacterium blinds and lyses a Chlamydomonad alga. Proc. Natl. Acad. Sci. U.S.A. 118, e2107695118.

Imai, S.I., Guarente, L., 2016. It Takes Two to Tango: Nad+ and Sirtuins in Aging/Longevity Control. npj Aging Mech. Dis. 2, 16017.

Itoh, E., Odakura, R., Oinuma, K.I., Shimizu, M., Masuo, S., Takaya, N., 2017a. Sirtuin e Is a Fungal Global Transcriptional Regulator That Determines the Transition from the Primary Growth to the Stationary Phase. J. Biol. Chem. 292, 11043–11054.

Itoh, E., Shigemoto, R., Oinuma, K.I., Shimizu, M., Masuo, S., Takaya, N., 2017b. Sirtuin A Regulates Secondary Metabolite Production by Aspergillus Nidulans. J. Gen. Appl. Microbiol. 63, 228–235.

Kawauchi, M., Nishiura, M., Iwashita, K., 2013. Fungus-Specific Sirtuin HstD Coordinates Secondary Metabolism and Development through Control of LaeA. Eukaryot. Cell 12, 1087–1096.

Khaldi, N., Seifuddin, F.T., Turner, G., Haft, D., Nierman, W.C., Wolfe, K.H., Fedorova, N.D., 2010. SMURF: Genomic Mapping of Fungal Secondary Metabolite Clusters. Fungal Genet. Biol. 47, 736–741.

Kim, H.K., Lee, S., Jo, S.M., McCormick, S.P., Butchko, R.A.E., Proctor, R.H., Yun, S.H., 2013. Functional Roles of FgLaeA in Controlling Secondary Metabolism, Sexual Development, and Virulence in Fusarium Graminearum. PLoS One 8, e68441.

Kosugi, S., Hasebe, M., Tomita, M., Yanagawa, H., 2009. Systematic Identification of Cell Cycle-Dependent Yeast Nucleocytoplasmic Shuttling Proteins by Prediction of Composite Motifs. Proc. Natl. Acad. Sci. U. S. A. 106, 10171–10176.

Krespach, M.K.C., Stroe, M.C., Netzker, T., Rosin, M., Zehner, L.M., Komor, A.J., Beilmann, J.M., Krüger, T., Scherlach, K., Kniemeyer, O., Schroeckh, V., Hertweck, C., Brakhage, A.A., 2023. Streptomyces Polyketides Mediate Bacteria–Fungi Interactions across Soil Environments. Nat. Microbiol. 8, 1348–1361.

Lee, I., Oh, J.H., Keats Shwab, E., Dagenais, T.R.T., Andes, D., Keller, N.P., 2009. HdaA, a Class 2 Histone Deacetylase of Aspergillus Fumigatus, Affects Germination and Secondary Metabolite Production. Fungal Genet. Biol. 46, 782–790.

Martín, J.F., 2017. Key Role of LaeA and Velvet Complex Proteins on Expression of β-Lactam and PR-Toxin Genes in Penicillium Chrysogenum: Cross-Talk Regulation of Secondary Metabolite Pathways. J. Ind. Microbiol. Biotechnol. 44, 525–535.

Medema, M.H., Blin, K., Cimermancic, P., De Jager, V., Zakrzewski, P., Fischbach, M.A., Weber, T., Takano, E., Breitling, R., 2011. AntiSMASH: Rapid Identification, Annotation and Analysis of Secondary Metabolite Biosynthesis Gene Clusters in Bacterial and Fungal Genome Sequences. Nucleic Acids Res. 39, 339–346.

Michishita, E., McCord, R.A., Boxer, L.D., Barber, M.F., Hong, T., Gozani, O., Chua, K.F., 2009. Cell Cycle-Dependent Deacetylation of Telomeric Histone H3 Lysine K56 by Human SIRT6. Cell Cycle 8, 2664–2666.

Min, J., Landry, J., Sternglanz, R., Xu, R.M., 2001. Crystal Structure of a SIR2 Homolog-NAD Complex. Cell 105, 269–279.

Misslinger, M., Scheven, M.T., Hortschansky, P., López-Berges, M.S., Heiss, K., Beckmann, N., Heigl, T., Hermann, M., Krüger, T., Kniemeyer, O., Brakhage, A.A., Haas, H., 2019. The Monothiol Glutaredoxin GrxD Is Essential for Sensing Iron Starvation in Aspergillus Fumigatus, PLoS Genetics.

Miyamoto, A., Kadooka, C., Mori, K., Tagawa, Y., Okutsu, K., Yoshizaki, Y., Takamine, K., Goto, M., Tamaki, H., Futagami, T., 2020. Sirtuin SirD Is Involved in α-Amylase Activity and Citric Acid Production in Aspergillus Luchuensis Mut. Kawachii during a Solid-State Fermentation Process. J. Biosci. Bioeng. 129, 454–466.

Nayak, T., Szewczyk, E., Oakley, C.E., Osmani, A., Ukil, L., Murray, S.L., Hynes, M.J., Osmani, S.A., Oakley, B.R., 2006. A Versatile and Efficient Gene-Targeting System for Aspergillus Nidulans. Genetics 172, 1557–1566.

Netzker T., Fischer J., Weber J., Mattern D.J., König C.C., Valiante V., Schroeckh V., Brakhage A.A., 2015 Microbial communication leading to the activation of silent fungal secondary metabolite gene clusters. Front. Microbiol. 6, 299.

Netzker, T., Flak, M., Krespach, M.K., Stroe, M.C., Weber, J., Schroeckh, V., Brakhage, A.A., 2018. Microbial Interactions Trigger the Production of Antibiotics. Curr. Opin. Microbiol. 45, 117–123.

Nützmann, H.W., Fischer, J., Scherlach, K., Hertweck, C., Brakhagea, A.A., 2013. Distinct Amino Acids of Histone H3 Control Secondary Metabolism in Aspergillus Nidulans. Appl. Environ. Microbiol. 79, 6102–6109.

Nützmann, H.W., Reyes-Dominguez, Y., Scherlach, K., Schroeckh, V., Horn, F., Gacek, A., Schümann, J., Hertweck, C., Strauss, J., Brakhage, A.A., 2011. Bacteria-Induced Natural Product Formation in the Fungus Aspergillus Nidulans Requires Saga/Ada-Mediated Histone Acetylation. Proc. Natl. Acad. Sci. U. S. A. 108, 14282–14287.

Oxford, A.E., Raistrick, H., Simonart, P., 1939. Studies in the Biochemistry of Micro-Organisms. Biochem. J. 33, 240–248.

Palmer, J.M., Bok, J.W., Lee, S., Dagenais, T.R.T., Andes, D.R., Kontoyiannis, D.P., Keller, N.P., 2013. Loss of CclA, Required for Histone 3 Lysine 4 Methylation, Decreases Growth but Increases Secondary Metabolite Production in Aspergillus Fumigatus. PeerJ 1, e4.

Palmer, J.M., Keller, N.P., 2010. Secondary Metabolism in Fungi: Does Chromosomal Location Matter? Curr. Opin. Microbiol. 13, 431–436.

Pidroni, A., Faber, B., Brosch, G., Bauer, I., Graessle, S., 2018. A Class 1 Histone Deacetylase as Major Regulator of Secondary Metabolite Production in Aspergillus Nidulans. Front. Microbiol. 9, 2212.

Romsdahl, J., Wang, C.C.C., 2019. Recent Advances in the Genome Mining of: Aspergillus Secondary Metabolites (Covering 2012-2018). Medchemcomm 10, 840–866.

Rösler, S.M., Kramer, K., Finkemeier, I., Humpf, H.U., Tudzynski, B., 2016. The SAGA Complex in the Rice Pathogen Fusarium Fujikuroi: Structure and Functional Characterization. Mol. Microbiol. 102, 951–974.

Sauve, A.A., Celic, I., Avalos, J., Deng, H., Boeke, J.D., Schramm, V.L., 2001. Chemistry of Gene Silencing: The Mechanism of NAD+-Dependent Deacetylation Reactions. Biochemistry 40, 15456–15463.

Schroeckh, V., Scherlach, K., Nutzmann, H.-W., Shelest, E., Schmidt-Heck, W., Schuemann, J., Martin, K., Hertweck, C., Brakhage, A.A., 2009. Intimate Bacterial-Fungal Interaction Triggers Biosynthesis of Archetypal Polyketides in Aspergillus Nidulans. Proc. Natl. Acad. Sci. 106, 14558–14563.

Shwab, E.K., Jin, W.B., Tribus, M., Galehr, J., Graessle, S., Keller, N.P., 2007. Historie Deacetylase Activity Regulates Chemical Diversity in Aspergillus. Eukaryot. Cell 6, 1656–1664.

Smith, K.M., Kothe, G.O., Matsen, C.B., Khlafallah, T.K., Adhvaryu, K.K., Hemphill, M., Freitag, M., Motamedi, M.R., Selker, E.U., 2008. The Fungus Neurospora Crassa Displays Telomeric Silencing Mediated by Multiple Sirtuins and by Methylation of Histone H3 Lysine 9. Epigenetics Chromatin 1, 5.

Stevenson, J.S., Liu, H., 2011. Regulation of White and Opaque Cell-Type Formation in Candida Albicans by Rtt109 and Hst3. Mol. Microbiol. 81, 1078–1091.

Tamura, K., Stecher, G., Peterson, D., Filipski, A., Kumar, S., 2013. MEGA6: Molecular Evolutionary Genetics Analysis Version 6.0. Mol. Biol. Evol. 30, 2725–2729.

Tennen, R.I., Bua, D.J., Wright, W.E., Chua, K.F., 2011. SIRT6 Is Required for Maintenance of Telomere Position Effect in Human Cells. Nat. Commun. 2, 433.

Tribus, M., Bauer, I., Galehr, J., Rieser, G., Trojer, P., Brosch, G., Loidl, P., Haas, H., Graessle, S., 2010. A Novel Motif in Fungal Class 1 Histone Deacetylases Is Essential for Growth and Development of Aspergillus. Mol. Biol. Cell 21, 345–353.

Troejer, P., Brandtner, E.M., Brosch, G., Loidl, P., Galehr, J., Linzmaier, R., Haas, H., Mair, K., Tribus, M., Graessle, S., 2003. Histone Deacetylases in Fungi: Novel Members, New Facts. Nucleic Acids Res. 31, 3971–3981.

Williams, R.B., Henrikson, J.C., Hoover, A.R., Lee, A.E., Cichewicz, R.H., 2008. Epigenetic Remodeling of the Fungal Secondary Metabolome. Org. Biomol. Chem. 6, 1895–1997.

Workman, J.L., Kingston, R.E., 1998. Alteration of Nucleosome Structure as a Mechanism of Transcriptional Regulation. Annu. Rev. Biochem. 67, 545–579.

Wurtele, H., Tsao, S., Lépine, G., Mullick, A., Tremblay, J., Drogaris, P., Lee, E.H., Thibault, P., Verreault, A., Raymond, M., 2010. Modulation of Histone H3 Lysine 56 Acetylation as an Antifungal Therapeutic Strategy. Nat. Med. 16, 774–780.

Yang, B., Miller, A., Kirchmaier, A.L., 2008. HST3/HST4-Dependent Deacetylation of Lysine 56 of Histone H3 in Silent Chromatin. Mol. Biol. Cell 19, 4993–5005.

Yu, J.H., Keller, N., 2005. Regulation of Secondary Metabolism in Filamentous Fungi. Annu. Rev. Phytopathol. 43, 437–458.

Zadra, I., Abt, B., Parson, W., Haas, H., 2000. XylP Promoter-Based Expression System and Its Use for Antisense Downregulation of the Penicillium Chrysogenum Nitrogen Regulator NRE. Appl. Environ. Microbiol. 66, 4810–4816.

Zhang, Y., Fan, J., Ye, J., Lu, L., 2020. The Fungal-Specific Histone Acetyltransferase Rtt109 Regulates Development, DNA Damage Response, and Virulence in Aspergillus Fumigatus. Mol. Microbiol. 109, 1–16.

Zocher, R., Paul, E., Madry, N., Peeters, H., Kleinkauf, H., Keller, U., Nihira, T., 1986. Biosynthesis of Cyclosporin A: Partial Purification and Properties of a Multifunctional Enzyme from Tolypocladium Inflatum. Biochemistry 25, 550–553.

