## Supplementary material for "Bacteria-induced activation of a fungal silent gene cluster is controlled by histone deacetylase Sirtuin E": Suplements

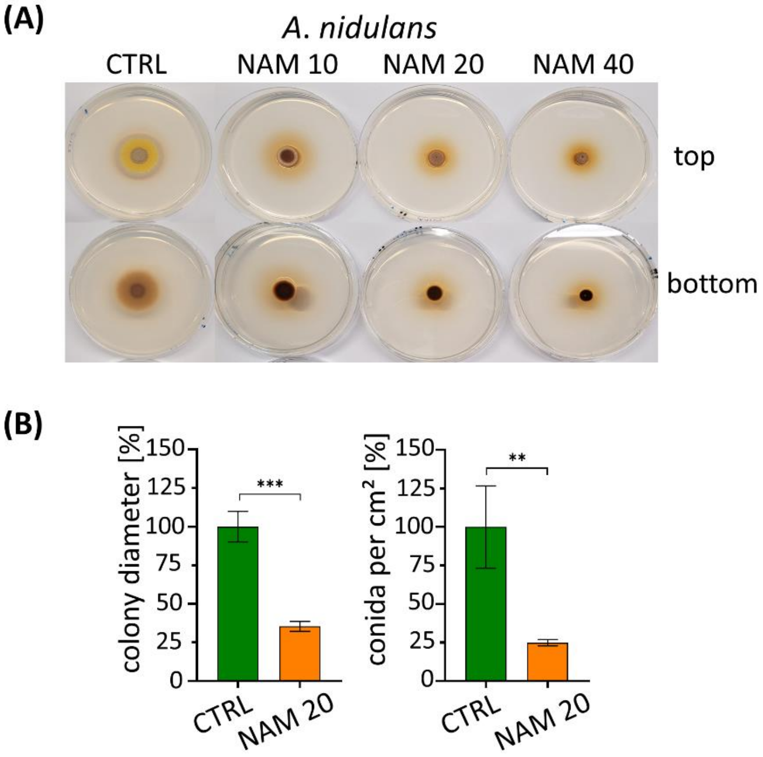

**Figure S1. Phenotypic analysis of *A. nidulans* wild type treated with NAM. (A)** *A. nidulans* colonies on AMM agar plates grown with or without 10-40 mM NAM from $10^6$ conidia each, cultivated for 3 d at 37°C. **(B)** Quantification of 20 mM NAM on colony diameter (n=3, bars represent mean ± SD). Significance was assessed by an unpaired *t*-test (** $p < 0.02$; *** $p < 0.002$).

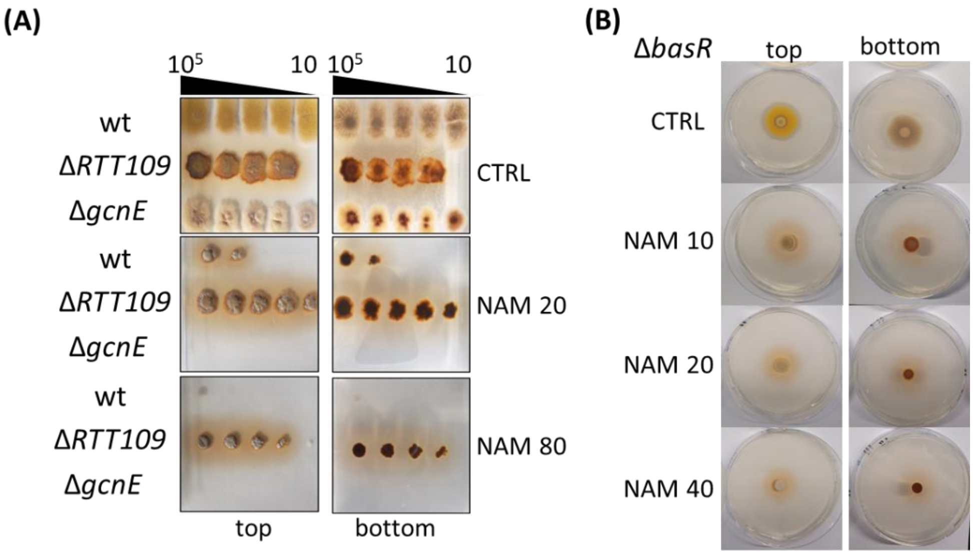

**Figure S2. Effects of NAM treatment on *A. nidulans* mutants with deleted *RTT109*, *gcnE* and *basR* gene. (A) Loss of *RTT109* leads to tolerance against high NAM concentrations.** Conidia ($10^5$–$10^1$) of *A. nidulans* wt, Δ*RTT109*, and Δ*gcnE* mutant strains were point-inoculated on AMM agar plates with NAM 20 mM and 80 mM or without (CTRL). Strains were cultivated for 5 d at 37 °C. Agar plates are shown from top and bottom. **(B) Loss of *basR* did not rescue the morphological and physiological defects caused by NAM treatment.** Conidia ($10^6$) of *A. nidulans* Δ*basR* strain were point-inoculated on AMM agar plates with 10, 20 and 40 mM NAM or without (CTRL). Strains were cultivated for 3 d at 37 °C.

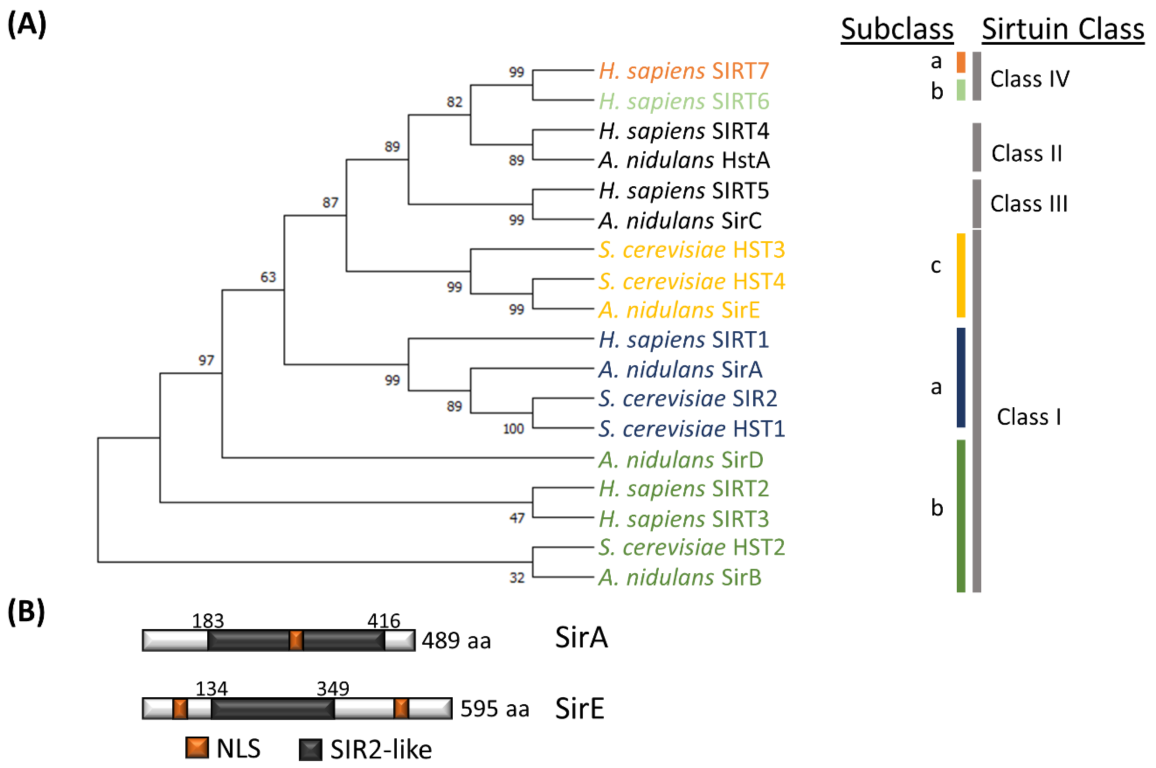

**Figure S3. Phylogenetic analysis of sirtuins of *S. cerevisiae, H. sapiens* and *A. nidulans* based on amino acid sequences. (A)** For phylogenetic analysis, MUSCLE alignment and neighbor joining were conducted using MEGA6 with 1000 bootstrap replicates on *A. nidulans*, *H. sapiens*, and *S. cerevisiae*. **(B)** Analysis of SirA and SirE *via* NLS mapper revealed the presence of putative nuclear localization sequences (NLSs) (Kosugi *et al.*, 2009) in addition to the SIR2-like domain.

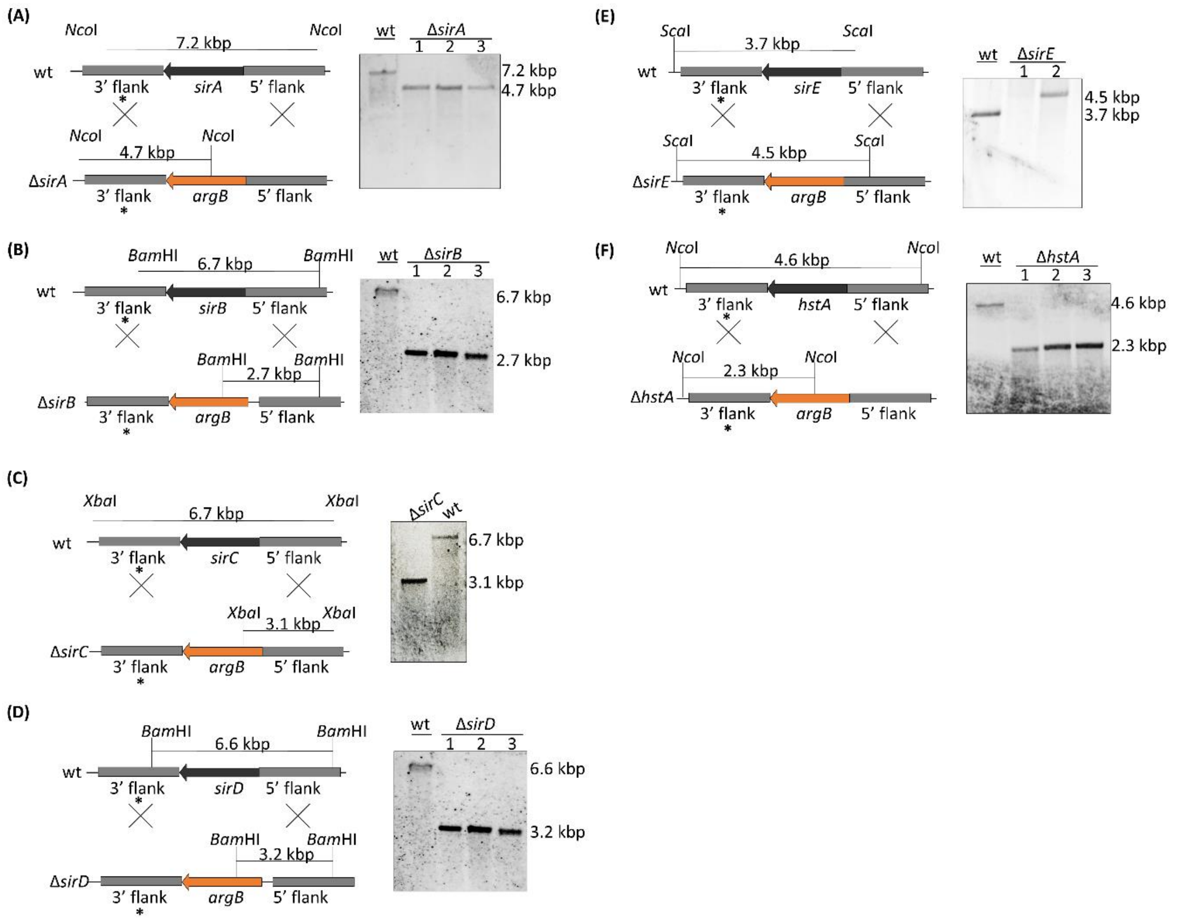

**Figure S4. Genomic organization and Southern blot analyses of sirtuin deletion mutants compared with wt. (A-F)** All putative sirtuin-encoding genes of *A. nidulans* were replaced by the *argB* gene *via* double homologous recombination. Transformants were selected for arginine prototrophy and checked by Southern blot analysis with a probe (*) directed against the flanking regions of sirtuin genes or *argB*, as indicated. Wt served as a control.

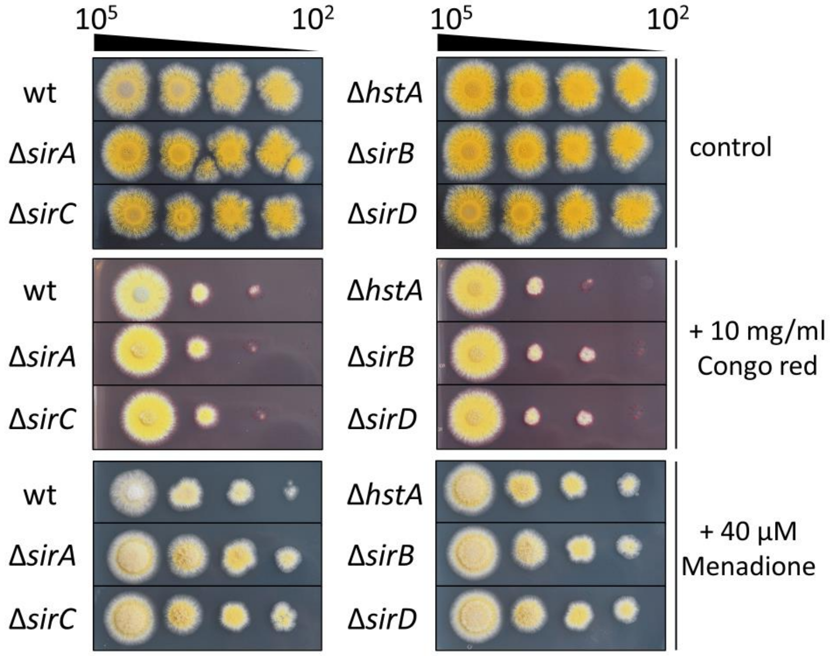

**Figure S5. Colonies of sirtuin deletion mutants in comparison to wt.** Conidia ($10^5$ to $10^2$) of *A. nidulans* wt, Δ*sirA*, Δ*sirB*, Δ*sirC*, Δ*sirD*, and Δ*hstA* strains were point-inoculated on AMM agar plates supplemented with Congo red (10 mg/ml) or with menadione (40 µM), and cultivated for 3 d at 37 °C.

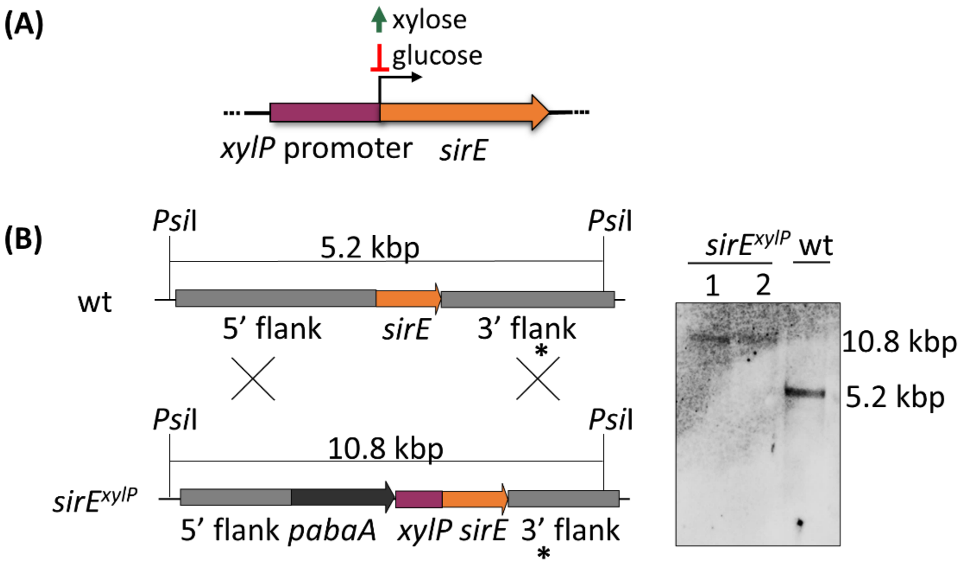

**Figure S6. Exchange of the native *sirE* promoter against the xylose-inducible *xylP* promoter (*xylP*p) from *P. chrysogenum*. (A)** While glucose represses the *xylP* promoter, the addition of xylose to the medium induces the expression of *xylP*p-controlled genes. **(B)** Schematic depiction of the *sirE* locus of the *A. nidulans* wt recipient strain and the *xylP*p-*sirE* strain (left) confirmed by Southern blot analysis (right). The DNA region to which the probe binds, is indicated by *.

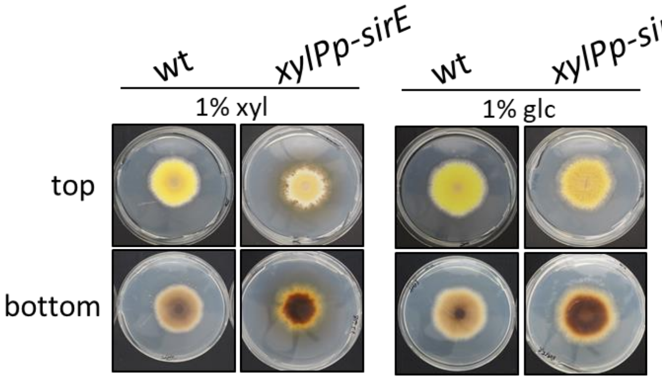

**Figure S7. Overexpression of *sirE* affects colony morphology.** Conidia ($10^5$) of wt and *xylP*p-*sirE* were point-inoculated on AMM agar plates, containing either 1% (w/v) glucose (glc) or 1% (w/v) xylose (xyl) and were cultivated for 3 d at 37 °C. Agar plates are shown from top and bottom.

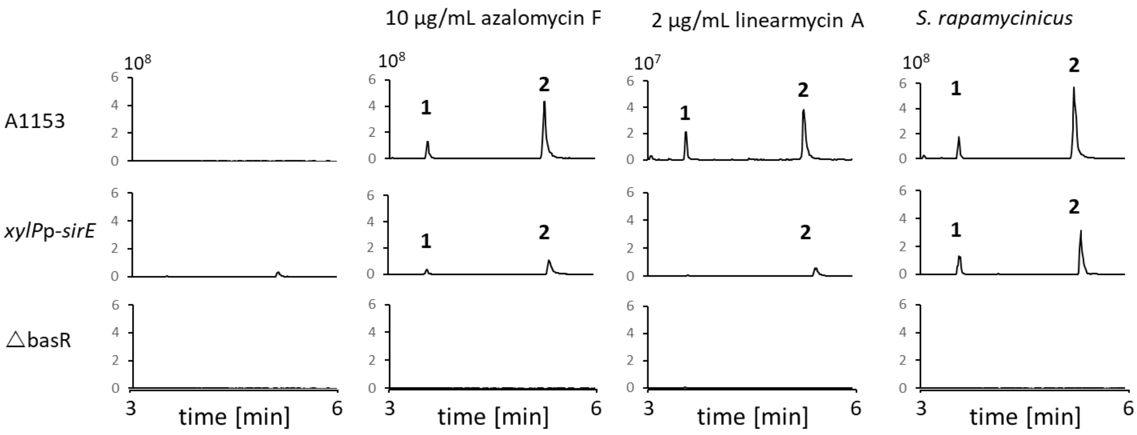

**Figure S8. Cocultures of *A. nidulans* strains with arginoketides and *S. rapamycinicus.*** LC-MS profiles of EICs are depicted for m/z 167 $[M-H]^-$. Marked peaks correspond to orsellinate (1) and lecanorate (2).

A

| Protein in *A. mulundensis* | Protein in *A. nidulans* | Function | Identity |
|---|---|---|---|
| DSM5745_09158 | BasR (AN7174) | Myb-like TF | 71.52% |
| DSM5745_06405 | SirE (AN1226) | $NAD^+$ dependent HDAC | 80.83% |
| DSM5745_00010 | OrsA (AN7909) | Iterative type I PKS | 80.83% |
| DSM5745_00008 | OrsB (AN7911) | Putative amidohydrolase | 86.36% |
| n.a. | OrsC (AN7912) | Putative tyrosinase | n.a. |
| DSM5745_00006 | OrsD (AN7913) | Unknown function | 68.07% |
| DSM5745_00005 | OrsE (AN7914) | Putative dehydrogenase | 80.00% |

B

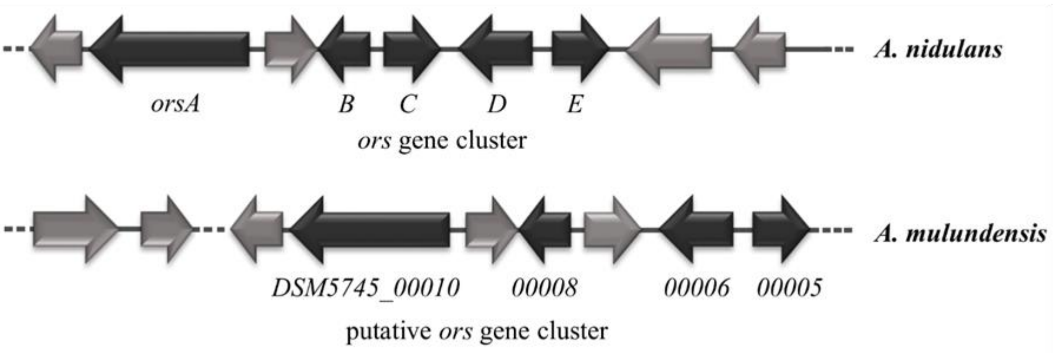

**Figure S9. Comparison of *A. nidulans ors* cluster with the putative *ors* cluster of *A. mulundensis.* (A)** BLASTp results of selected proteins linked to *ors* gene cluster activation in *A. nidulans* and the corresponding homologs of *A. mulundensis* DSMZ 5745. **(B)** Schematic alignment of the putative *ors* gene cluster of *A. mulundensis* with the *ors* gene cluster of *A. nidulans*. According to antiSMASH and BLASTp analyses, the core genes of the putative *ors* gene cluster of *A. mulundensis* is composed of DSM5747_00010 (*orsA*), DSM5747_00008 (*orsB*), DSM5747_00006 (*orsD*), and DSM5747_00005 (*orsE*). A homologous gene of *orsC* (AN7912), encoding a putative tyrosinase, is missing.

### Materials and methods

**Microorganisms, media and cultivation.** All microorganisms used in this study are listed in Table S1. If not otherwise stated, *A. nidulans* strains were grown in *Aspergillus* minimal medium (AMM) (Brakhage and Van den Brulle, 1995) at 37 °C with 200 rpm. When required, AMM was supplemented with L-arginine (871 µg/mL), *p*-aminobenzoic acid (3 µg/mL) and pyridoxine HCl (5 µg/mL). Pre-cultures were inoculated with 4 × $10^8$ spores per ml. Samples for RNA extraction or HPLC analysis were taken after 24 h, if not stated otherwise. For repressing conditions, the *xylP*p-*sirE* strain was grown in presence of 1 % (w/v) glucose as the sole carbon source, while 1 % (w/v) xylose was used for inducing conditions. NAM (Roth) was freshly prepared and dissolved in water and sterile filtered. *A. mulundensis* was grown on malt agar (20 g $L^{-1}$ malt extract, 2 g $L^{-1}$ yeast extract, 10 g $L^{-1}$ glucose, 0,25 g $L^{-1}$ $NH_4Cl$, 0,25 g $L^{-1}$ $K_2HPO_4$) or AMM for 1-10 d at 26 °C.

**Generation of sirtuin deletion strains.** All plasmids in this study were assembled according to the Gibson assembly method (Gibson *et al.*, 2009). In brief, the NEBuilder® HiFi DNA assembly Master mix (New England Biolabs, Frankfurt a. M., Germany) was employed according to the manufacturer's instructions. The *sirE* deletion strain was generated by amplifying the 5'-flanking region with primer pairs NJ24/NJ25, while primer pairs NJ22/NJ43 were used to amplify the 3'-flank of *sirE* from genomic DNA. Primers NJ24 and NJ43 contained 30-bp complementary overhangs to the 5'- and 3'-ends of the *argB* cassette, respectively. Primer pairs NJ63/NJ64 were used to amplify the pJET1 vector with complementary 30-bp ends to the 5'- and 3'-flanks of *sirE.* The *argB* cassette was amplified using the primer pair argcassfor/argcassrev from plasmid pLUC-T*argB*02 as a template. The linear DNA fragments were seamlessly assembled according to the manufacturer's protocol. The resulting plasmid pJETΔ*sirE* was linearized with *Afl*II and *Psi*I and used to transform *A*. *nidulans* wt strain A1153. Putative clones were analyzed by colony PCR and correct genomic reorganization verified by Southern hybridization (Figure S4C). Plasmids containing the deletion constructs for the remaining sirtuins *hstA*, *sirA*, *sirB*, *sirC*, and *sirD* were also generated by seamless assembly of PCR-amplified DNA fragments (see Table S2) as described above in detail for Δ*sirE*. The assembled plasmid pJET Δ*sirB* was linearized with *Nde*I and *Pvu*II; pJET Δ*hstA* was linearized with *Cla*I; pJET Δ*sirC* was linearized with *Nhe*I and *Cla*I; pJET Δ*sirD* was linearized with *Cla*I; and the Δ*sirA* cassette was PCR-amplified from pJET Δ*sirA* with primer pair NJ34/NJ50. Deletion cassettes were purified and *A*. *nidulans* A1153 was transformed with the cassettes. Putative clones were analyzed by colony PCR and by Southern hybridization (Figure S4A-F).

**Generation of the *A. nidulans xylP*p-*sirE* strain.** Primer pairs NJ189/NJ225 and NJ186/NJ224 were used to PCR-amplify the 5'- and 3'-flanking regions of *sirE* from *A*. *nidulans* wt genomic DNA, respectively. The *pabaA* cassette was amplified with the primer pair pabacassfor/NJ223 from plasmid pAbaAnid. Primer NJ223 contained a 25-bp complementary overhang to the 5'-end of the *xylP* promoter, while NJ225 contained a 25-bp complementary overhang to the 5'-end of the *pabaA* cassette. The *xylP* promoter was amplified by PCR with the primer pair NJ213/NJ214 from plasmid pUC18 *xylP*p-*npgA*. Primer NJ213 contained a 25-bp complementary end to the 5'-end of *sirE*. Primer pair NJ197/NJ198 was used to amplify the pJET1 vector backbone with complementary 30-bp ends to the 5'- and 3'-flanks of *sirE.* The linear DNA fragments were seamlessly assembled according to the manufacturer's instructions. *A*. *nidulans* wt was transformed with the resulting plasmid pJET *pabaA*-*xylP*p-*sirE* linearized with *Dra*III and *Psi*I. Putative clones were analyzed by PCR and correct genomic reorganization was verified by Southern blot analysis (Figure S4B).

**Preparation of chromosomal DNA and Southern blot analysis.** *A. nidulans* genomic DNA was isolated as previously described (Schroeckh *et al*., 2009). Southern blotting was performed using a digoxigenin-11-dUTP-labeled probe (Jena Bioscience, Jena, Germany), as previously described (Schroeckh *et al*., 2009). Primers for amplification of probes are listed in Table S2.

**qRT-PCR.** Expression levels were quantified by qRT-PCR according to the ΔΔCt method as previously described (Schroeckh *et al*., 2009). Total RNA was purified with the Universal RNA Purification Kit (roboklon, Berlin, Germany) according to the manufacturer's instructions. Reverse transcription was performed with RevertAid Reverse Transcriptase (Thermo Fisher Scientific, Darmstadt, Germany) for 3 h at 46 °C. Quantitative RT-PCR of 5 ng cDNA samples was performed with MyTaq HS mix 2x (Bioline, Luckenwalde, Germany) and EvaGreen (Biotium, Fremont, California, USA) using primers for the respective target genes (Table S3). The *A. nidulans* ß-actin gene (*AN6542*) was used as an internal standard for calculation of mRNA steady-state levels, as previously described (Schroeckh *et al*., 2009).

**Extraction of NPs and HPLC analysis.** Liquid chromatography- mass spectrometry (LC-MS) analyses were performed as described (Fischer *et al*., 2018). Briefly, whole fungal cultures were homogenized using a T25 ULTRA-TURRAX® (IKA, Staufen, Germany) homogenizer. The homogenized cultures were extracted twice with an equal volume of ethyl acetate. Two spatula of anhydrous sodium sulfate were added to remove remaining water. Then, the samples were filtered in pointed pistons and transferred to a Laborota 4000 efficient rotary evaporator (Heidolph Instruments, Schwabach, Germany). The dried extracts were dissolved in 1 mL methanol and filtered through a 0.2 µm polytetrafluoroethylene filter (Carl Roth GmbH, Karlsruhe). Afterwards, the samples were analyzed by LC-$MS^2$ on an UltiMate™ 3000 binary rapid-separation liquid chromatograph with photodiode array detector (PDA) and an LTQ XL™ linear ion trap mass spectrometer (Thermo Fisher Scientific, Dreieich, Germany) equipped with an electrospray ion source. The extracts were analyzed on a 150 mm- by 4.6-mm Accucore™ RP-MS LC column with a particle size of 2.6 µm (Thermo Fisher Scientific, Dreieich, Germany) at a flow rate of 1 mL $min^{-1}$, with 0.1 % (v/v) HCOOH-MeCN/0.1 % (v/v) and the following gradient over 21 min: initial 0 % (vol/vol) acetonitrile increased to 80 % (v/v) over 15 min and then to 100 % (v/v) over 2 min, held at 100 % (v/v) for 2 min and reversed to 0 % (v/v) over 2 min) The injection volume amounted to 10 µL.

For figure S8, the samples were analyzed by LC-MS on a Vanquish Flex UHPLC system equipped with a PDA and an Orbitrap$^{TM}$ Exploris 120 mass spectrometer (Thermo Fisher Scientific, Dreieich, Germany). The extracts were separated on a LUNA$^{R}$ Omega PS C18 LC-column with a particle size of 1.6 µm (Phenomenex, Aschaffenburg, Germany) at a flow rate of 0.5 mL $min^{-1}$ with the same solvents and the following gradient over 13 min: initial 5 % (v/v) acetonitrile hold for 0.5 min and increased to 100 % (v/v) over 8 min held at 100 % (v/v) for another 2 minutes and reversed to 5 % (v/v) over 2 min. The injection volume amounted to 2 µL.

**Phenotypic analysis.** Conidia of *A. nidulans* strains were diluted in droplet solution (0.9 % (w/v) NaCl, 0.01 % (v/v) Tween-20, 0.1 % (w/v) agarose) and point-inoculated on AMM or malt agar plates.

**Phylogenetic and statistical analysis.** The Molecular Evolutionary Genetics Analysis version 6 (MEGA6) software was employed for phylogenetic analyses of SirE orthologs (Tamura *et* *al*., 2013). Deduced amino acid sequences were obtained from UniProtKB database (http://www.uniprot.org). Statistical analysis was conducted using the GraphPad Prism version 6.0 (GraphPad Software Inc., San Diego, USA).

**Table S1: Microorganisms used in this study.**

| Name | Genotype | Reference |
|---|---|---|
| *Aspergillus nidulans* | | |
| A1153 | *yA1*, *pabaA1*; *argB2*; *pyroA4*, *nkuA*::*bar* | (Nayak *et al*., 2006) |
| Δ*basR* | A1153 *basR*::*argB* | (Fischer *et al*., 2018) |
| Δ*RTT109* | A1153 *AN8807*::*argB* | (Nützmann *et al*., 2011) |
| Δ*gcnE* | A1153 *gcnE*::*argB* | (Nützmann *et al*., 2011) |
| Δ*sirE* | A1153 *sirE*::*argB* | This study |
| Δ*sirB* | A1153 *sirB*:*argB* | This study |
| Δ*hstA* | A1153 *hstA*::*argB* | This study |
| Δ*sirC* | A1153 *sirC*::*argB* | This study |
| Δ*sirD* | A1153 *sirD*::*argB* | This study |
| Δ*sirA* | A1153 *sirA*::*argB* | This study |
| *xylP*p-*sirE* | A1153 *sirE*::*pabaA*-*xylP*p-*sirE* | This study |
| *orsA*p-*GFP*-*nluc* | RMS011 *orsA*::*GFP*::*nluc*::*pabaA* | (Krespach *et al*., 2023) |
| *A. mulundensis* | | |
| *CBS 140610* | Wt | Westerdijk fungal biodiversity institute |
| *Escherichia coli* | | |
| XL1-Blue | *recA1*, *endA1*, *gyrA96*, *thi-1*, *hsdR17*, *supE44*, *relA1*, *lac* [F´ *proAB lacI*$^q$ *Z*Δ*M15* *Tn10* (*Tet*$^R$ )] | Stratagene |

**Table S2: Strategies for the generation of sirtuin-knockout plasmids by Gibson assembly.**

| Plasmid name | DNA fragments | Size (kb) | Template DNA | Primer name |
|---|---|---|---|---|
| pJET | 5'-flank | 1.0 | gDNA A1153 | NJ30/NJ31 |
| Δ*sirB* | 3'-flank | 1.0 | gDNA A1153 | NJ32/NJ33 |
| | pJET1 | 3.2 | pJET1 | NJ67/NJ68 |
| | *argB* cassette | 2.7 | pLUC-T*argB02* | argcassfor/argcassrev |
| pJET | 5'-flank | 1.0 | gDNA A1153 | NJ28/NJ29 |
| Δ*hstA* | 3'-flank | 1.0 | gDNA A1153 | NJ26/NJ27 |
| | pJET1 | 3.2 | pJET1 | NJ71/NJ72 |
| | *argB* cassette | 2.7 | pLUC-T*argB02* | argcassfor/argcassrev |
| pJET | 5'-flank | 1.0 | gDNA A1153 | NJ36/NJ50 |
| Δ*sirA* | 3'-flank | 1.0 | gDNA A1153 | NJ34/NJ35 |
| | pJET1 | 3.2 | pJET1 | NJ69/NJ70 |
| | *argB* cassette | 2.7 | pLUC-T*argB02* | argcassfor/argcassrev |
| pJET | 5'-flank | 1.0 | gDNA A1153 | NJ14/NJ47 |
| Δ*sirC* | 3'-flank | 1.0 | gDNA A1153 | NJ16/NJ17 |
| | pJET1 | 3.2 | pJET1 | NJ65/NJ66 |
| | *argB* cassette | 2.7 | pLUC-T*argB02* | argcassfor/argcassrev |
| pJET | 5'-flank | 1.0 | gDNA A1153 | NJ18/NJ19 |
| Δ*sirD* | 3'-flank | 1.0 | gDNA A1153 | NJ20/NJ21 |
| | pJET1 | 3.2 | pJET1 | NJ73/NJ74 |
| | *argB* cassette | 2.7 | pLUC-T*argB02* | argcassfor/argcassrev |

**Table S3: List of oligonucleotides.**

Complementary overhangs are marked in bold, inserted restriction sites in italics and underlined.

| Designation | Sequence (5' - 3') | Target |
|---|---|---|
| qacnfwd | CACCCTTGTTCTTGTTTTGCTC | *AN6542* fw. (*actA*) |
| Qacnrev | AAGTTCGCTTTGGCAACGC | *AN6542* rev. (*actA*) |
| qorsAfor | CTATACCACCGATAGCCAGGAC | *AN7909* fw. (*orsA*) |
| qorsArev | CAGTGAGCAGGGCAAAGAAG | *AN7909* rev. (*orsA*) |
| qorsDfor | GCAACGAGCCTGACATTACC | *AN7913* fw. (*orsD*) |
| qorsDrev | CCGCACATCAACCATCTCTG | *AN7913* rev. (*orsD*) |
| pabacassfor | TGCCAGATCTGTAGAAAGGTC | *pabaA* cassette fw. |
| argcassfor | ATGGGAGTCAAAGTTCTGTTTGC | *argB* cassette fw. |
| argcassrev | GGAAGCGAGAGAACATGTCAA | *argB* cassette rev. |
| NJ14 | TACCGATCTGCCATCCTCCA | *sirC* 5'-flank fw. |
| NJ16 | **AATAACTAATTGACATGTTCTCTCGCTTCC**TAATGCTACCGTACTAAATGTTGG | *sirC* 3'-flank fw. |
| NJ17 | CCTGCATTGTTGACGAGTAC | *sirC* 3'-flank rev. |
| NJ18 | AAATTCGTGGTTTTATTGGCTAAA | *sirD* 3'-flank fw. |
| NJ19 | **AATAACTAATTGACATGTTCTCTCGCTTCC**ATGCCCATATTTTCACTTAGCA | *sirD* 3'-flank rev. |
| NJ20 | **GATCAGGGCAAACAGAACTTTGACTCCCAT**TTTCTGTTGGAGAAAAGAGTGT | *sirD* 5'-flank fw. |

| | | |
|---|---|---|
| NJ21 | GAACGCTTCTGGGCTGTA | *sirD* 5'-flank rev. |
| NJ22 | CGCCGACTTATTTAATACTCCTT | *sirE* 5'-flank fw. |
| NJ24 | **AATAACTAATTGACATGTTCTCTCGCTTCC**AGGATTCACTATTTTTGATATTATT | *sirE* 3'-flank fw. |
| NJ26 | AGGGCTCGACGAAGGGGGTT | *hstA* 5'-flank fw. |
| NJ27 | **GATCAGGGCAAACAGAACTTTGACTCCCAT**CGGAGTTGTCGCCGCTGG | *hstA* 5'-flank rev. |
| NJ28 | **AATAACTAATTGACATGTTCTCTCGCTTCC**ATTTATACATTTAAACAGTAGAGTA | *hstA* 3'-flank fw. |
| NJ29 | TCCGAAATCACCCAGCTT | *hstA* 3'-flank rev. |
| NJ30 | CTGGTTAACTTTCGTGACTG | *sirB* 3'-flank fw. |
| NJ31 | **AATAACTAATTGACATGTTCTCTCGCTTCC**AACCGCAAAGATCTAAAGA | *sirB* 3'-flank rev. |
| NJ32 | **GATCAGGGCAAACAGAACTTTGACTCCCAT**TGGCGTGCTGGTGTCTGT | *sirB* 5'-flank fw. |
| NJ33 | GAACCATTTCTGCCGCAAA | *sirB* 5'-flank rev. |
| NJ34 | ACTGGATGATCTCTGTCTTTTG | *sirA* 3'-flank fw. |
| NJ35 | **AATAACTAATTGACATGTTCTCTCGCTTCC**CATAGCGGTGGAGTTTCTGG | *sirA* 3'-flank rev. |
| NJ36 | **GATCAGGGCAAACAGAACTTTGACTCCCAT**CCTGGTGATGGGATTGAG | *sirA* 5'-flank fw. |
| NJ43 | **GATCAGGGCAAACAGAACTTTGACTCCCAT**GGATGACGCGAGAGTAGAGTAC | *sirE* 5'-flank rev. |
| NJ47 | **GATCAGGGCAAACAGAACTTTGACTCCCAT**GGTGAGCAGAAAACGGGT | *sirC* 5'-flank rev. |
| NJ50 | CAGATCTTTCGTAGGTCGCC | *sirA* 5'-flank rev. |
| NJ56 | GCGGGTACATGCCACAATAC | *basR* fw. qRT |
| NJ57 | TCTCGGGCATCATCAACTCC | *basR* rev. qRT |
| NJ63 | **TAAAGTAAGGAGTATTAAAATAAGTCGGCG**TAGAAGATCTCCTACAATATTCTCAGC | pJET rev. (*sirE*) |
| NJ64 | **GATAACACGTACTGAGAAGACAGATAAGAA***GCTAGC*AACTCGAGCCATCCGGAT | pJET fw. (*sirE*) |
| NJ65 | **TGCGTAGCCGTGGAGGATGGCAGATCGGTA**TAGAAGATCTCCTACAATATTCTCAGC | pJET rev. (*sirC*) |
| NJ66 | **GCGAATTGATGTACTCGTCAACAATGCAGG***TTTAAA*AACTCGAGCCATCCGGAT | pJET fw. (*sirC*) |
| NJ67 | **TGCCGGAATTCAGTCACGAAAGTTAACCAG**TAGAAGATCTCCTACAATATTCTCAGC | pJET rev. (*sirB*) |
| NJ68 | **TTCAGGGCGCCTTTGCGGCAGAAATGGTTC***CATATG*AACTCGAGCCATCCGGAT | pJET fw. (*sirB*) |
| NJ69 | **GCTCGGCCCAAAAGACAGAGATCATCCAGT**TAGAAGATCTCCTACAATATTCTCAGC | pJET rev. (*sirA*) |
| NJ70 | **AAAAAACAGGGGCGACCTACGAAAGATCTG***ATCGAT*AACTCGAGCCATCCGGAT | pJET fw. (*sirA*) |
| NJ71 | **TCTCCCCCGCAACCCCCTTCGTCGAGCCCT**TAGAAGATCTCCTACAATATTCTCAGC | pJET rev. (*hstA*) |
| NJ72 | **AAAGGGCAATTAAAGCTGGGTGATTTCGGA***ATCGAT*AACTCGAGCCATCCGGAT | pJET fw. (*hstA*) |
| NJ73 | **TAAGAATTTAGCCAATAAAACCACGAATTT**TAGAAGATCTCCTACAATATTCTCAGC | pJET rev. (*sirD*) |
| NJ74 | **CTGGTCGCCTTCTACAGCCCAGAAGCGTTC***ATCGAT*AACTCGAGCCATCCGGAT | pJET fw. (*sirD*) |
| NJ186 | ACGGTGTTTGTTTCTTATCTGTC | *sirE* 3'-flank rev. |

| | | |
|---|---|---|
| NJ189 | CAGAAGCCAAGTAGCATTCC | *sirE* 5’-flank fw. |
| NJ197 | **TATGATCCTAGGAATGCTACTTGGCTTCTG**TAGAAGATCTCC TACAATATTCTCAGC | pJET fw. (*sirE*) |
| NJ198 | **TAATCCTTGATGCATTGATTAGCATTCGCC**AACTCGAGCCAT CCGGAT | pJET rev. (*sirE*) |
| NJ213 | **CGGGCTTGGTTTTGCGAGGTGCCAT**GGTTGGTTCTTCGAGT CGATG | *xylP* rev. (*sirE*) |
| NJ214 | ACTGATGCGAGCAACAGTATG | *xylP* fw. (*sirE*) |
| NJ216 | GCGCACAAAGAAAGAGCCAG | *sirE* fw. qRT |
| NJ217 | GCATTGCCTTATGCTCCTCC | *sirE* rev. qRT |
| NJ223 | **CTGGCATACTGTTGCTCGCATCAGT**ATCTGGACATGCGACG GAG | *pabaA* rev. (*sirE*) |
| NJ224 | ATGGCACCTCGCAAAACC | *sirE* gene fw. |
| NJ225 | **CGCAGACCTTTCTACAGATCTGGCA**GGATGACGCGAGAGTA GAGTAC | *sirE* 5’-flank rev. |
